# Expansion of gamma-butyrolactone signaling molecule biosynthesis to phosphotriester natural products

**DOI:** 10.1101/2020.10.11.335315

**Authors:** Yuta Kudo, Takayoshi Awakawa, Yi-Ling Du, Peter A. Jordan, Kaitlin E. Creamer, Paul R. Jensen, Roger G. Linington, Katherine S. Ryan, Bradley S. Moore

## Abstract

Bacterial hormones, such as the iconic gamma-butyrolactone A-factor, are essential signaling molecules that regulate diverse physiological processes, including specialized metabolism. These low molecular weight compounds are common in *Streptomyces* species and display species-specific structural differences. Recently, unusual gamma-butyrolactone natural products called salinipostins were isolated from the marine actinomycete genus *Salinispora* based on their anti-malarial properties. As the salinipostins possess a rare phosphotriester motif of unknown biosynthetic origin, we set out to explore its construction by the widely conserved 9-gene *spt* operon in *Salinispora* species. We show through a series of in vivo and in vitro studies that the *spt* gene cluster dually encodes the saliniphostins and newly identified A-factor-like gamma-butyrolactones (Sal-GBLs). Remarkably, homologous biosynthetic gene clusters are widely distributed amongst many actinomycete genera, including *Streptomyces,* suggesting the significance of this operon in bacteria.

## Introduction

Actinobacteria are a rich source of specialized metabolites that have been developed into life-saving drugs. Recent advances in genome sequencing and mining have revealed that Actinobacteria have far greater potential for secondary metabolite production than previously realized.^1^ Yet, much of this potential remains cryptic, as many biosynthetic genes are poorly expressed in normal laboratory incubation conditions.^2^ The manipulation of the signaling mechanisms for gene expression can be a key to activate the expression of dormant biosynthetic genes.^3–5^ However, the regulation of biosynthetic pathways, and the autoregulators themselves, remains poorly understood.

Among the known signaling molecules, gamma-butyrolactones (GBLs) are recognized to be involved in the regulation of morphological development and secondary metabolism in actinomycete bacteria, especially in the genus of *Streptomyces*.^6–10^ In contrast, among the metabolically prolific marine actinomycete genus *Salinispora*, far less is known about the signaling pathways regulating the biosynthesis of their rich repertoire of natural products.^11–15^ However the recently reported potent and selective anti-malarial compounds, salinipostins (Figure 1A, salinipostin A (**1**)) from *Salinispora* sp. RL08-036-SPS-B,^16,17^ may shed some light on this. Salinipostins possesses a GBL ring analogous to A-factor, as well as a highly unusual phosphotriester ring. Two compounds, cyclipostins and cyclophostin (Figure 1C), from *Streptomyces* have this same structural motif and have been reported as hormone sensitive lipase and acetylcholinesterase inhibitors, respectively.^18,19^ These compounds are estimated to be derived from a similar pathway as the GBL biosynthetic pathway based on their structural similarity.

**Figure 1.**
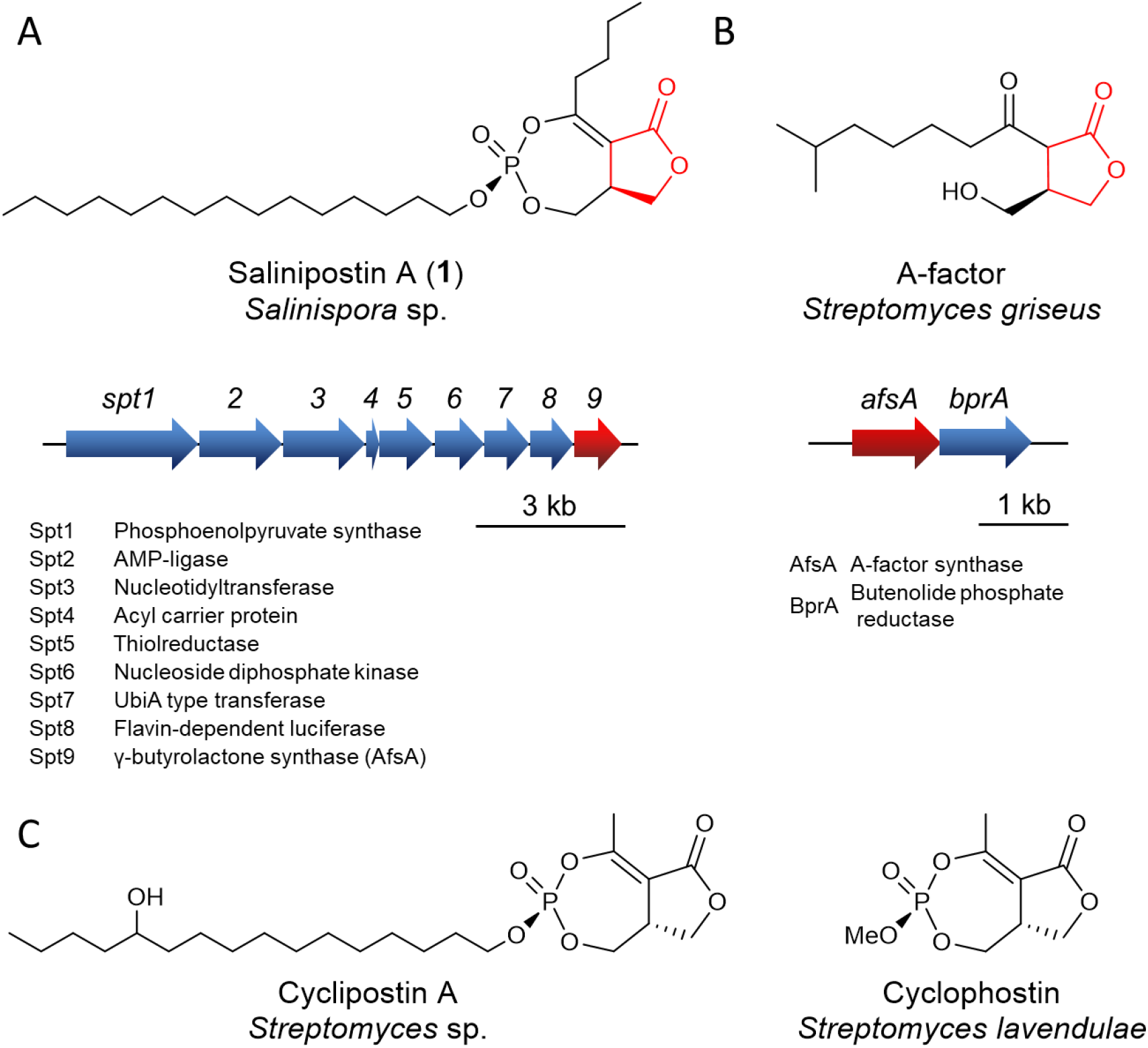
Chemical structures of natural phosphotriester compounds and A-factor with their sources and biosynthetic gene clusters. (A) Salinipostin A and its biosynthesis gene cluster from *Salinispora tropica* CNB-440^2^; (B) A-factor and its biosynthesis gene cluster^8,22^; (C) cyclipostin A and cyclophostin, whose biosynthetic genes have not yet been identified/predicted^18,19^. Gamma-butyrolactone structure and key relevant biosynthetic gene were highlighted in red in panel (A) and (B).

The most studied GBL compound is A-factor (Figure 1B) from *Streptomyces griseus*.^20,21^ The GBL structure is formed from the condensation of beta-ketoacyl acyl carrier protein (ACP) and dihydroxyacetone phosphate (Figure 1B) by the A-factor synthase AfsA followed by successive reduction by BprA and dephosphorylation.^22^ AfsA homologues are common in streptomycetes where they have been shown to be involved not just in GBL biosynthesis but also in the production of gamma-butenolides and furans.^6,23–26^

Recently, we showed that the AfsA homologue Spt9 in *Salinispora* bacteria is responsible for the construction of the salinipostins and its volatile byproducts, salinilactones.^2,27,28^ The *spt9* gene is the terminal gene of a nine-gene biosynthetic locus (Figure 1A) that is broadly conserved in *Salinispora* species^29^ (*spt* was annotated as *butyrolactone 1* in this reference). Herein, we report the functional characterization of *spt* genes, *spt6* and *spt9*, toward salinipostin biosynthesis and show that the *spt* gene cluster is also responsible for the synthesis of novel, natural GBLs as co-products of salinipostin biosynthesis.

## Results and discussion

### Discovery of gamma-butyrolactone compounds from *Salinispora* species

We previously reported the global transcriptional activity of biosynthesis gene clusters in *Salinispora* species and showed that the *spt* locus was involved with salinipostin biosynthesis through genetic inactivation of *spt9* and concomitant loss of salinipostin.^2^ Upon re-examination of the *S. tropica* CNB-440 *spt9*-deletion mutant in comparison with the native CNB-440 strain by Liquid Chromatography-Mass Spectrometry (LC-MS), we identified compounds in addition to the salinipostins that were also abolished in the mutant (Figure 2B). Inspection of the molecular formulae of compounds **2**(C_11_H_19_O_4_, *m/z* 215.1280, [M+H]^+^ calcd. 215.1278) and **3**(C_10_H_17_O_4_, *m/z* 201.1126, [M+H]^+^ calcd. 201.1121) suggested that they could be simple, A-factor-like GBL compounds (Figure S1).

**Figure 2.**
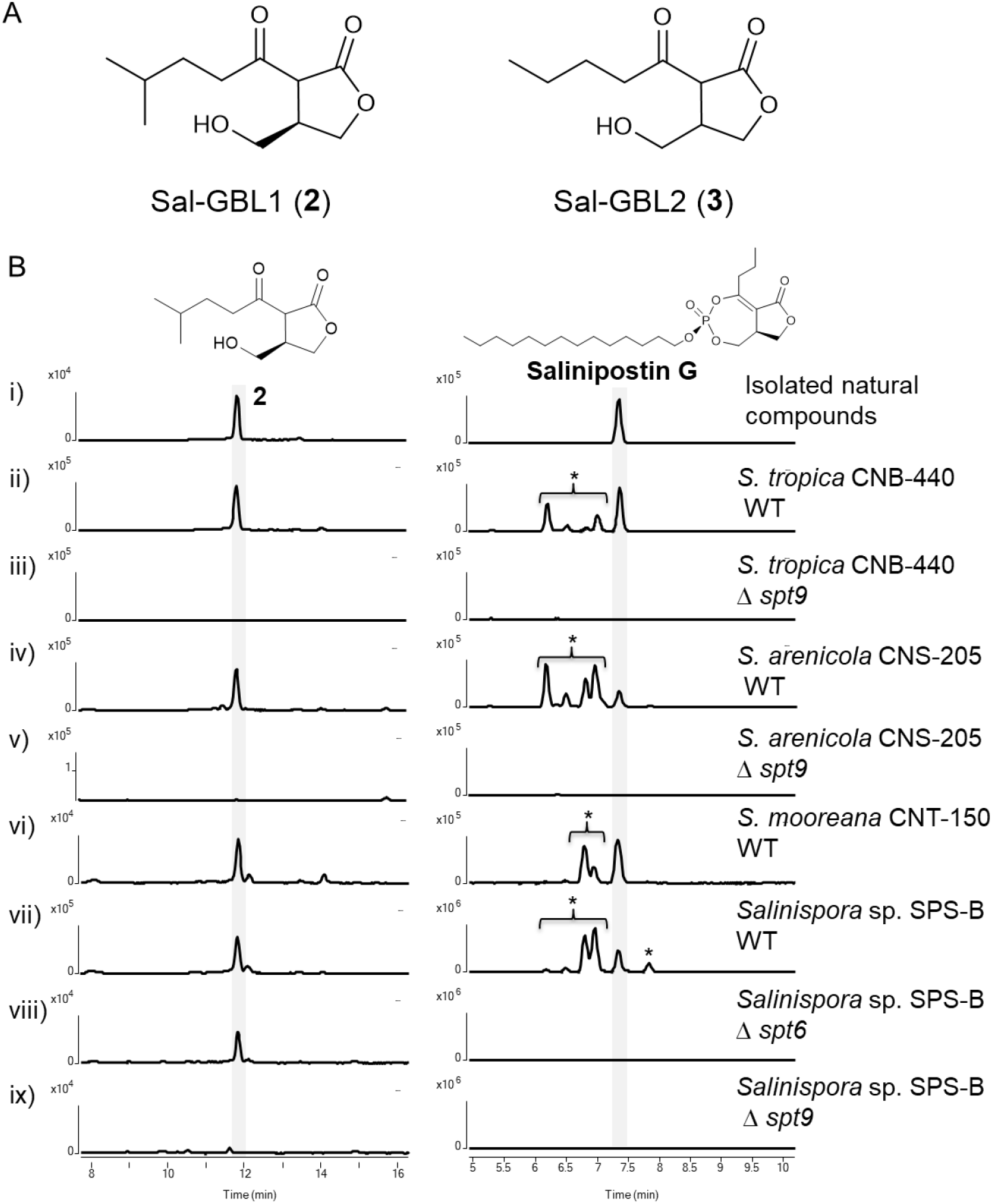
Structures and mutagenesis of *spt* products. (A) Structures of new natural GBLs isolated from *S. tropica* CNB-440, Sal-GBL1 (**2**) and Sal-GBL2 (**3**); (B) EIC at *m/z* 215.1278 (Sal-GBL1, left column) and *m/z* 445.2714 (salinipostin G, right column) for i) isolated natural compounds, ii) *S. tropica* CNB-440 wild type, iii) *S. tropica* CNB-440 Δ*spt9*, iv) *S. arenicola* CNS-205 wild type, v) *S. arenicola* CNS-205 Δ*spt9*, vi) *S. mooreana* CNT-150, vii) *Salinispora* sp. RL08-036-SPS-B wild type, viii) *Salinispora* sp. RL08-036-SPS-B Δ*spt6*, and ix) *Salinispora* sp. RL08-036-SPS-B Δ*spt9*. *Estimated as salinipostin analogues based on the high-resolution MS and/or MS/MS fragmentation pattern (Figures S4-8).

Generally, the production levels of GBLs in native strains are quite modest, reflecting the low effective concentration of the signaling molecules (e.g., A-factor, 10^−9^ M)^30^. As such, significant effort is often required to isolate and characterize GBLs as in the case of SCB-1 and IM-2 in which several hundred liters of cultures were analyzed.^31,32^ To enhance the poor productivity of the *S. tropica* GBL compounds, we grew strain CNB-440 with Amberlite XAD-7HP resin to achieve an approximately 50-fold enhancement of GBL production (Figure S2). With this method, we purified 1.2 mg of the most abundant GBL (Sal-GBL1, **2**) from 3 L culture for comprehensive NMR analysis (Figures S20–23). Based on the 2D NMR analysis, the structure of **2** was confirmed as a new natural GBL containing an iso-hexanoyl sidechain (Figure 2A). The circular dichroism spectrum of **2** exhibited similar Cotton effects as the natural form of synthetic 3-(*R*)-A-factor^33,34^ (Figure S24), therefore establishing the stereochemistry at C-3 in **2** as (*R*). This stereochemical configuration is consistent with the stereochemistry at C-3 in salinipostin^16^. A second analogue (Sal-GBL2, **3**) was elucidated with a pentanoyl sidechain (Figure 2A, **3**) based on the LC-MS comparison with a chemically synthesized standard (Figure S3, 25, 26).

We observed that compounds **2**and **3** are also produced in the other species of *Salinispora* producing salinipostins, namely *S. arenicola* CNS-205, *S. mooreana* CNT-150 and *Salinispora* sp. RL08-036-SPS-B (Figure 2B, Figure S3), representing the first identification of A-factor-like GBL compounds in the genus of *Salinispora*. The production of **2**, **3**, and salinipostins were also completely abolished in *S. arenicola* CNS-205 Δ*spt9* and *Salinispora* sp. RL08-036-SPS-B Δ*spt9*, reinforcing that *spt9* is essential for their biosynthesis (Figure 2B, Figure S3).

On the basis of the distinct chemical features of the salinipostins and Sal-GBLs and the gene composition of the *spt* gene cluster, we suspected that the biosynthetic pathway likely diverges at an early common intermediate. As such, we knocked out the putative nucleotide diphosphate-kinase *spt6* gene in *Salinispora* sp. RL08-036-SPS-B and showed that this mutant produced **2**, which lack a phosphotriester, but not salinipostins, with the phosphotriester group (Figure 2B). These results are consistent with our proposed, diverging pathway (Figure 3), whereby subsequent pyrophosphate formation via Spt6 redirects the biosynthetic pathway to salinipostins.

**Figure 3.**
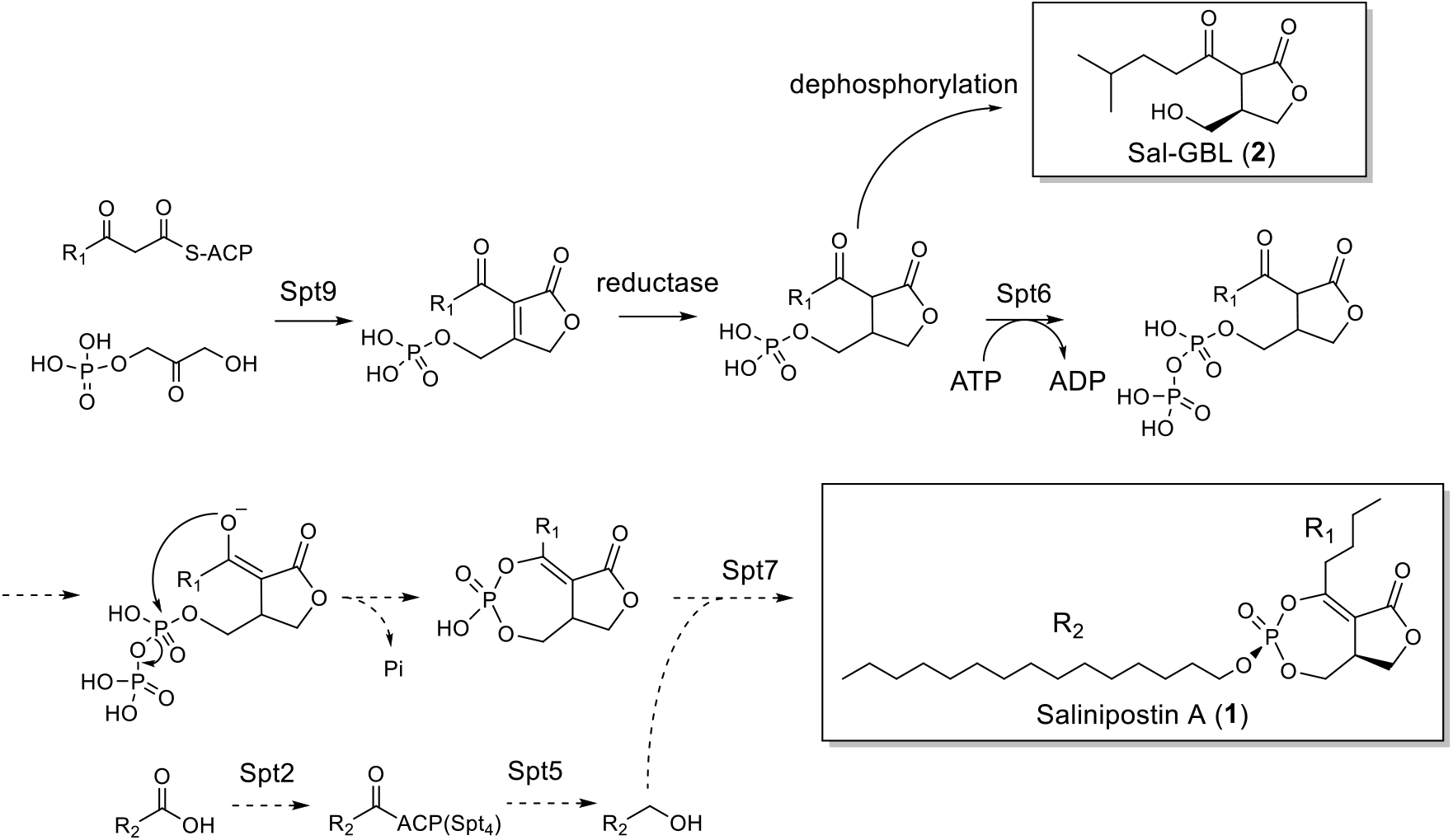
Enzymatic pathway toward Sal-GBL (**2**) and Sal-GBL pyrophosphate and hypothetical biosynthetic pathway to salinipostin A (**1**). Solid arrows represent confirmed biochemical reactions, while dashed arrows are proposed. See Figure 1A for the putative functions of the Spt proteins.

### *In vitro* reconstitution of the gamma-butyrolactone structure

To further evaluate salinipostin biosynthesis and its branchpoint with GBL synthesis, we next evaluated the *in vitro* activity of recombinant Spt9 based on prior examination of the homologous A-factor synthase AfsA^22^. We prepared the N-terminus maltose-binding protein (MBP) and histidine tagged Spt9 (Spt9-H-MBP) from *E. coli* harboring pET28a-MBP-*spt9* and removed both affinity tags using TEV protease (Figure S18). We next synthesized both substrates: the beta-keto heptanoyl *N*-acetylcysteamine thioester (SNAC) (compound **4**) as a substrate mimic of the ACP-bound substrate and dihydroxyacetone (**5**).^35^ We first observed the consumption of **4** in the enzymatic reaction mixture, however, a product peak was not detected in the LC-MS analysis. Therefore, we analyzed the reaction mixture using LC-MS immediately after short incubation periods. Incubation of tag-free Spt9 with substrates **4** and **5** (1 mM, each) in a phosphate-citrate buffer at 30 °C for 10 min yielded a plausible phosphorylated butenolide compound (**7**) having the anticipated molecular formula C_10_H_17_O_7_P (*m/z* 277.0481, [M-H]^−^ calcd. 277.0483) and phosphate fragment ion at *m/z* 96.9709 ([M-H]^−^ calcd. for H_2_O_4_P, 96.9696) (Figure 4B, Figure S15).

**Figure 4.**
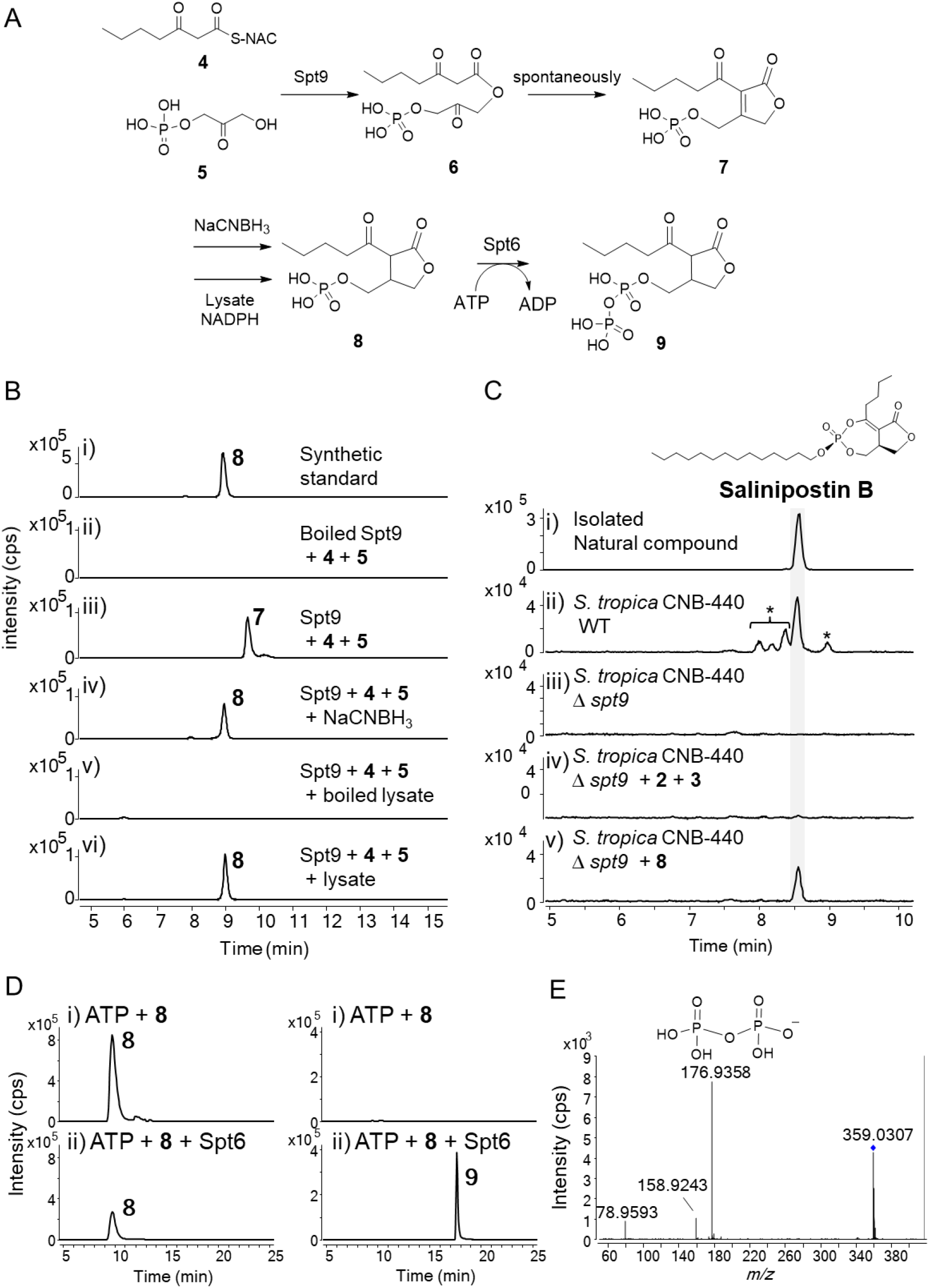
Gamma-butyrolactone structure formation in the biosynthesis of salinispostin. (A) *In vitro* reaction scheme, (B) Extracted ion chromatograms (EICs) for *in vitro* reaction mixtures: (i) Synthetic standard **8**, (ii) **4** and **5** treated with boiled Spt9, (iii) **4** and **5** treated with Spt9, (iv) **4** and **5** treated with Spt9, then with NaCNBH_3_, (v) **4** and **5** treated with Spt9, then with boiled lysate, and (vi) **4**and **5**treated with Spt9, then with lysate. EIC at *m/z* 279.0639 for (i), (iv), (v) and (vi), and *m/z* 277.0483 for (ii) and (iii). (C) EICs at *m/z* 459.2870 for XAD extracts from the feeding experiment: (i) isolated salinipostin B, (ii) *S. tropica* CNB-440 wild-type, (iii) *S. tropica* CNB-440 Δ*spt9,* (iv) *S. tropica* CNB-440 Δ*spt9* supplemented with **2**and **3**, and (v) *S. tropica* CNB-440 Δ*spt9* supplemented with **8**. *Estimated as the salinipostin analogues based on the high-resolution MS and MS/MS fragmentation pattern (Figures S9,10). (D) HILIC-LC-MS EICs for *in vitro* reaction of phosphorylation of Sal-GBL2 phosphate (**8**), (i) **8**, ATP and Mg^2+^ without H-Spt6 (ii) **8**, ATP and Mg^2+^ with H-Spt6. EICs at *m/z* 279.0639 (**8**, left column) and *m/z* 359.0302 (**9**, right column): (E) MS/MS spectrum of **9** with the structure of fragment ion, *m/z* 177.

In A-factor biosynthesis, butenolide formation is followed by reduction with BprA, which is encoded immediately downstream of *afsA* ^22^. The *spt* gene cluster, however, does not encode a *bprA* homologue nor elsewhere nearby. Thus, we chose to incubate enzymatically produced **7** with the lysate prepared from the *S. tropica* CNB-440 Δ*spt9* deletion mutant along with NADPH (Figure 4B). In this way we produced compound **8**(*m/z* 279.0632, ([M-H]^−^ calcd. 279.0639), which we validated by LC-MS analysis with synthetically prepared GBL-phosphate (**8**). Chemical reduction of the C-C double bond in **7** with NaBH3CN also gave a relatively stable compound that eluted at the same retention time as the synthetic GBL-phosphate (**8**). These results are consistent with previous reports where the expression of recombinant AfsA in *E. coli* resulted in the production of A-factor like compounds in the host and reduction of butenolide was suggested to be catalyzed by the endogenous reductase in the bacteria.^8,22^ Thus, despite their low sequence homology at 35.8% (pairwise positive, ClustalW alignment), our data unequivocally confirms that Spt9, like AfsA, catalyzes butenolide formation.

Although we assumed that passive cellular uptake of GBL-phosphate is unlikely based upon its polarity and negative charge, supplementation of *S. tropica* CNB-440 Δ*spt9* with synthetic **8** recovered the production of salinipostin B, which contains *n*-butyl side chain (Figure 4C, S13). Conversely, the supplementation of GBLs **2** and **3** did not recover salinipostin production. Only in the case of supplementation with **8** were other possible salinipostin analogues containing the C4H8-sidechain also detected (Figures S11, S12, S14), while natural salinipostins containing different sidechains were not generated (Figure 4C, (ii),(v)). These results support our hypothesis that salinipostin biosynthesis is initiated with the construction of the GBL. Furthermore, compound **3** was detected in the extract of *S. tropica* CNB-440 Δ*spt9* supplemented with synthetic **8**. Consistent with that observation, lysate of *S. tropica* CNB-440 Δ*spt9* similarly showed weak dephosphorylation activity. The incubation of **8** with commercial alkaline phosphatase (CIP) reliably gave **3** (Figure S16). Because there are no dedicated phosphatases in the *spt* gene cluster, we assume that this dephosphorylation was catalyzed by a nonspecific bacterial phosphatase, as similarly implied in A-factor biosynthesis.

Our investigations of salinipostin biosynthesis indicated the GBL-phosphate **8** is a biosynthetic intermediate of salinipostin. As a prelude to phosphodiester ring formation, we hypothesized that activation of the phosphate group of **8** is an absolute requirement. Biochemically this is achieved through pyrophosphate formation as seen in the formation of the phosphodiester ring in cyclic-AMP, where adenylate cyclase converts ATP into cyclic AMP along with releasing a pyrophosphate as a leaving group^36^. Our mutant analysis confirmed that the *spt6* gene encoding a diphosphate kinase is essential for the production of salinipostin as described above. Therefore, we further investigated the *in vitro* function of Spt6. We tested the enzymatic phosphorylation using the His-tagged Spt6 (H-Spt6) purified from *E. coli* harboring pET28a-*spt6* (Figure S19). Incubation of H-Spt6 and **8** with ATP and Mg^2+^ in HEPES buffer (pH 8) yielded the GBL-pyrophosphate (**9**), showing a fragment ion at *m/z* 176.9358 ([M-H]^−^ calcd. for H3O7P2, 176.9359) derived from the pyrophosphate group in the both hydrophilic interaction chromatography (HILIC) LC-MS/MS (Figure 4D, E) and reversed-phase (RP) LC-MS/MS analyses (Figure S17). Thus, we hypothesize that **9** is the first dedicated biosynthetic intermediate in the salinipostin pathway and is required for the formation of the cyclic phosphodiester group. Few enzymes catalyze phosphorylation reactions on phosphorylated substrates in secondary metabolism. One example is CalQ, which forms a diphosphate secondary metabolite, the protoxin phosphocalyculin A from toxic calyculin A^37^, but Spt6 shares low sequence identity with CalQ (13.9%, ClustalW alignment).

We anticipate that the remainder of the pathway from the GBL pyrophosphate **9** to the salinipostins and its highly unusual phosphotriester core involves additional enzymes encoded within the *spt* locus as proposed in Figure 3. Biochemical experiments are presently underway.

### Distribution and gene content of salinipostin-like biosynthetic gene clusters

The salinipostin biosynthetic gene cluster (*spt1-9*) is conserved at the genus level in *Salinispora*^29^. Using a targeted genome-mining approach, we identified nine genera in addition to *Salinispora* that maintain *spt*-like biosynthetic gene clusters (Figure 5). The *spt*-like gene clusters were identified in six diverse Actinobacterial families: *Nocardiaceae, Tsukamurellaceae, Mycobacteriaceae, Pseudonocardiaceae, Dietziaceae,* and *Streptomycetaceae*. Notably, *spt*-like gene clusters were found outside of the *Streptomycetaceae* where the gamma-butyrolactone A-factor and *spt9* homolog *afsA* were characterized. *Spt1, spt3*, *spt5*, *spt6*, *spt7*, and *spt9* are conserved across all *spt*-like gene clusters (Figure 5), although *spt2-3* and *spt6-9* gene fusions were observed in some strains. *Spt8* was uniquely observed in the *Salinispora spt* gene cluster. The organization of the *spt* biosynthetic gene cluster is largely conserved across multiple families of bacteria, ultimately suggesting that these diverse taxa have the capability to produce salinipostin-like phophotriester GBLs. A detailed analysis of *spt* gene distribution, organization, and evolutionary history will be the subject of another study (Creamer *et al*. 2020 bioRxiv).

**Figure 5.**
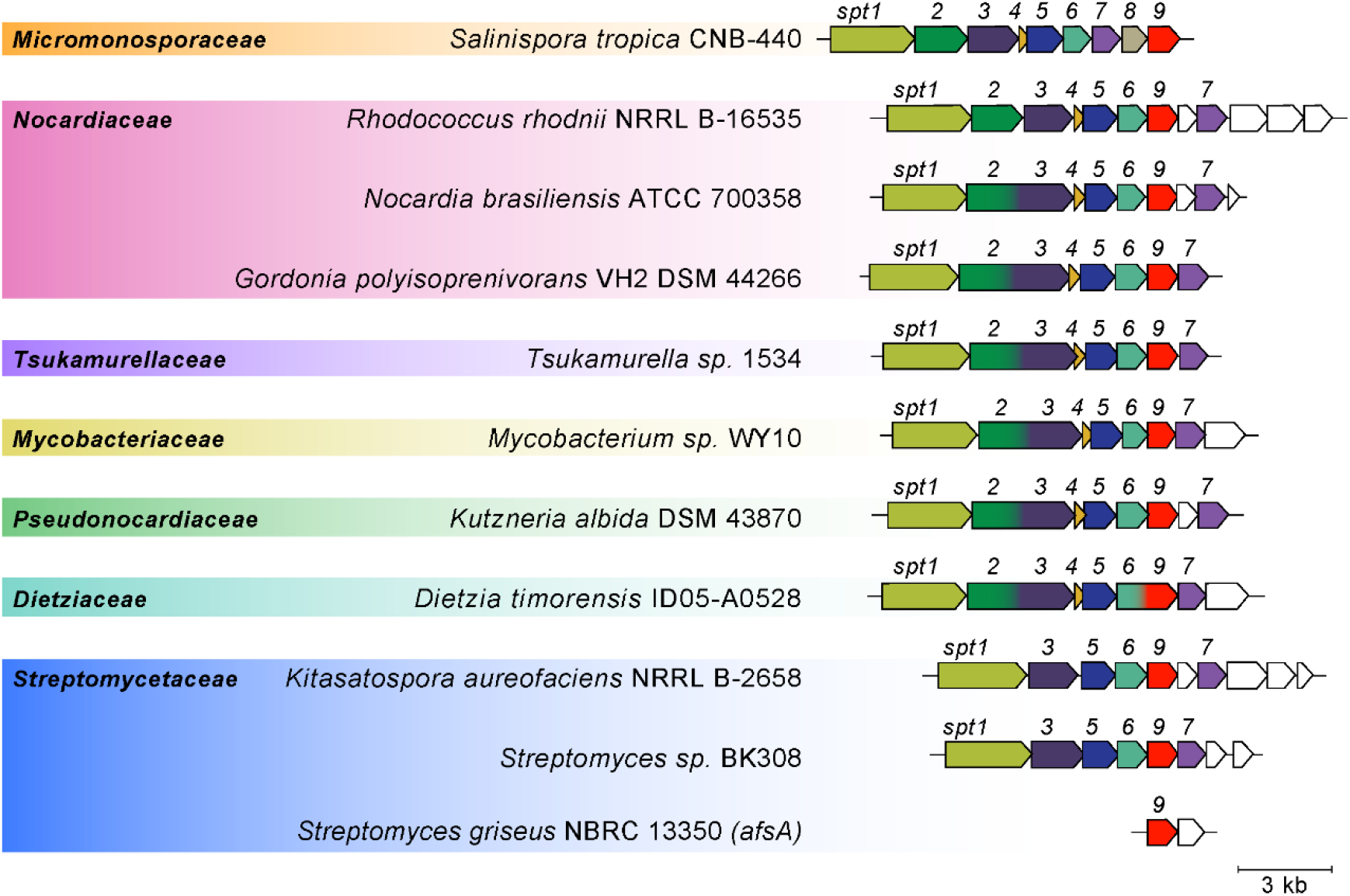
*Spt*-like biosynthetic gene clusters observed in diverse actinomycetes. Genes are colored by conserved Pfam function relative to *spt1-9* in *S. tropica*; white indicates no relation to *spt1-9*. Spt2-3 and Spt6-9 represent gene fusions. Products are known from *S. tropica* (salinipostins and Sal-GBLs) and *S. griseus* (A-factor).

In conclusion, we established the early steps of salinipostin biosynthesis in *Salinispora* bacteria and showed that the phosphorylated GBL **8** is the branchpoint metabolic intermediate to the salinipostins and the newly identified Sal-GBL1 (**2**) and Sal-GBL2 (**3**) A-factor-like compounds. Intriguingly, the *spt* locus may thus have a dual biological purpose in the construction of both butyrolactone chemotypes whose native *Salinispora* functions have yet to be determined. While no small molecules from *Salinispora* have yet been experimentally established as autoregulators, *Salinispora* also produce acyl-homoserine lactones that are known to mediate quorum sensing in gram-negative bacteria.^38^ Future investigations on the function of these molecules will undoubtedly shed light on the regulatory mechanism of secondary metabolism in *Salinispora*.

Our work also opens the door to establishing the biochemical logic for the construction of salinipostin’s highly unusual phosphotriester functionality that is shared by the streptomycete molecules cyclipostin and cyclophostin. The high energy GBL pyrophosphate **9** is chemically predisposed for an intramolecular cyclization reaction with loss of phosphate to a cyclic phosphodiester GBL intermediate as shown in Figure 3. Condensation with a fatty alcohol constructed with Spt2/4/5 theoretically completes the pathway to the salinipostins. This proposed pathway anticipates the biosynthesis of cyclipostin and cyclophostin, which together with the salinipostins, may be more common in actinomycete biology than previously thought given the wide distribution of *spt*-like biosynthesis gene clusters (Figure 5). As such, we hypothesize that salinipostin-like phosphotriester GBLs may function as a new class of signaling molecules. Interrogation of salinipostin biosynthetic gene clusters may represent a key step in further unlocking the biosynthetic potential in *Salinispora* and other actinomycetes.

## METHODS

### General

The high-resolution MS spectra were recorded on an Agilent 6530 Accurate-Mass qTOF spectrometer, equipped with a dual ESI ionization source and an Agilent 1260 LC system. Acquisition parameters of the mass spectrometer were following, range 80–850 *m/z*, MS scan rate 2/sec, MS/MS scan rate 3/sec, fixed collision energy 20 eV, source gas temperature 300°C, gas flow 11 L/min, nebulizer 45 psig, scan source parameters: VCap 3000, fragmentor 175 V, skimmer 65 V, Octopole RF Peak 750 V. LC-MS grade acetonitrile (MeCN), water and formic acid were purchased from Thermo Fisher Scientific (Waltham, Massachusetts, USA). Figures for mass spectral data were created in Mass Hunter (Agilent Technologies). Low-resolution LC-MS measurements were carried out on a Bruker Amazon SL ESI-Ion Trap mass spectrometer with Agilent Technologies 1200 Series HPLC system. LC-MS data was processed using Bruker Compass Data Analysis. The NMR spectra were recorded on a BRUKER Avance (600 MHz, CryoProbe) spectrometer in the solvents indicated. Signals are reported in ppm with the internal chloroform signal at 7.26 ppm (^1^H) and 77.16 ppm (^13^C), or the internal CD_3_OD signal at 3.31 ppm (^1^H) and 49.0 ppm (^13^C) as standard. Circular dichroism (CD) measurement of Sal-GBL1 (**2**) was obtained on an Aviv CD spectrometer model 62DS using a 1 nm bandwidth in a 0.5 cm cell at a concentration of 7.0 mM at 25°C, 3 sec/scan, 200 nm to 500 nm.

### Bacterial strains

*Salinispora tropica* CNB-440 (accession: NC_009380.1), *S. arenicola* CNS-205 (accession: NC_009953.1), *S. mooreana* CNT-150 (accession: NZ_AQZW00000000 or KB900614.1), *S. oceanensis* CNT-854 (HQ642928.1) and *Salinispora sp*. RL08-036-SPS-B (KP250536.1) were used in this study. These *Salinispora* strains were cultured on A1 agar plates (10 gL^−1^ starch, 4 gL^−1^ yeast extract, 2 gL^−1^ peptone, 18 gL^−1^ agar, 1 L artificial seawater). Starter cultures were inoculated from the colony grown on the A1 agar plate into 50 ml medium A1 (10 gL^−1^ starch, 4 gL^−1^ yeast, 2 gL^−1^ peptone, 1 L artificial seawater) in 250 ml Erlenmeyer flasks containing a stainless steel spring and grown for 3 or 4 days at 28°C with shaking at 230 RPM (New Brunswick Innova 2300).

### Gene deletion of *spt9*

#### i. Salinispora tropica

CNB-440/Δ*spt9* was generated in the previous study.^2^

#### ii. Salinispora arenicola

CNS-205/Δ*spt9*.

DNA fragment A containing the upstream 3.0 kb region of *spt9* was amplified by PCR with primer 1, 5’-cccccgggctgcaggaattc**acctcgaacgcgcctactggcac**-3’ (annealing sequence boldfaced) and 2, 5’-caccatcgatggtta**tagccgggtgtcggtcatcggttg**-3’, and fragment B containing the downstream 3.0 kb region of *spt9* was amplified by PCR with primer 3, 5’-accgacacccggcta**taaccatcgatggtgatctgtcgg**-3’ and 4, 5’-ccagcctacacatcgaattc**gcctgcgcaacgcatagtggagatc**-3’.Fragment A and B were assembled with pIJ773-derived fragment amplified by primer 5, 5’-gaattcctgcagcccgggggatc-3’ and 6, 5’-gaattcgatgtgtaggctggag-3’ using Gibson assembly technique (kit was purchased from New England Biolabs), to yield pIJ773-Δ*spt9*. The gene deletion vector pIJ773-Δ*spt9* was introduced into *S. arenicola* CNS-205 via intergeneric conjugation. The Apr^*R*^ exoconjugates from double-crossover were selected as previously reported procedure^39^. *S. arenicola* CNS-205/Δ*spt9* was isolated from Apr^*S*^ clones selected from the repeated subculture of Apr^*R*^ clones on the aparamycin-free agar plate.^27^

#### iii. *Salinispora* sp. RL08-036-SPS-B/Δ*spt9 and Salinispora* sp. RL08-036-SPS-B/Δ*spt6*

The gene inactivation experiments of *spt9* and *spt6* in *Salinispora* sp. RL08-036-SPS-B were performed using a PCR-targeting method. The coding region of *spt9* was first replaced with an apramycin resistant gene *aac(3)IV* in cosmid 12E4-bla, which was isolated from the genomic library of *Salinispora* sp. RL08-036-SPS-B. The *aac(3)IV* disruption cassette was amplified from pHY773 by PCR primers with homology extensions targeting *spt9* (primer pair, 5’-atgaccgacacccggctactgaccccacctcacacgacc**attccggggatccgtcgacc**-3’, and 5’-ctgagcctctcgtcgagcaccccggtcaacacctggcgt**tgtaggctggagctgcttc**-3’). The resulting vector 12E4-bla (Δ*spt9*) was passaged through *E. coli* S17-1 and then introduced into *Salinispora* sp. RL08-036-SPS-B via intergeneric conjugation. The apramycin^R^ exconjugants were screened by PCR for the double cross-over mutants. Inactivation of *spt6* was conducted similarly, except that an *aac(3)IV*-*ermE*\*p disruption cassette (primer pair, 5’-cctgacccacgatccggacaagcagcgctatcacggcac**tcagttcccgccagcctgct**-3’, and 5’-gaaagcgataccgcgtggctgcccaccacgatgaaatcc**taccaaccggcacgattgtg**-3’) was used to prevent the potential polar effects on downstream genes.

### LC-MS analysis of salinipostins and Sal-GBLs

Each strain was cultured in a 250 mL Erlenmeyer flask containing 50 mL of medium A1+BFe (A1 plus 1 gL^−1^ CaCO_3_, 40 mgL^−1^ Fe_2_(SO_4_)_3_‧4H_2_O, 100 mgL^−1^ KBr) at 28°C with shaking at 230 RPM (New Brunswick Innova 2300). Sterile Amberlite XAD-7HP resin (1 g, Sigma-Aldrich) was added to each flask after 48 hr, and the fermentation was continued for an additional 4 days. The resins and cells were collected by filtration using cheese cloth or centrifugation, washed with distilled water, and soaked in acetone for 2 hr. The extract was concentrated *in vacuo* and the resultant residue dissolved in methanol. After filtration, an aliquot of the extract was subjected to LC-MS. For the salinipostin analysis, HR-LC-MS analysis was performed using a Phenomenex Kinetex XB-C_18_ column (2.6 μm, 100 × 4.6 mm, 100 Å) with the following conditions; positive mode, mobile phase A: 0.1% formic acid in H_2_O, mobile phase B: 0.1% formic acid in MeCN, 1.0 mL/min, 0–8 min (80–90% B), 8-13 min (90%–100% B), 13-15 min (100% B). The ion at *m/z* 445.2714 (salinipostin G and analogues), 459.2870 (salinipostin B and analogues) and 473.3027 (salinipostin analogues) corresponding to the [M+H]^+^ ion was analyzed in the extracted ion chromatograms (EIC). Low-resolution LC-MS analysis was performed using a Phenomenex Kinetex XB-C_18_ column (2.6 μm, 100 × 4.6 mm, 100 Å) with the following conditions; positive mode, mobile phase A: 0.1% formic acid in H_2_O, mobile phase B: 0.1% formic acid in MeCN, 1.0 mL/min, 0–8 min (80–90% B), 8-10 min (90%–100% B), 10-15 min (100% B), with a flow splitter. For the Sal-GBLs, LC-MS analysis was performed using a Phenomenex Luna C18(2) column (5.0 μm, 150 x 4.6 mm, 100 Å) with the following conditions; positive mode, mobile phase A: 0.1% formic acid in H_2_O, mobile phase B: 0.1% formic acid in MeCN, 0.7 mL/min, 0–1 min (20% B), 1–21 min (20%–100% B), and 21–31 min (100% B). Ions at *m/z* 215.1278 and 201.1121 (Sal-GBL1 (**2**) and Sal-GBL2 (**3**), respectively) corresponding to the [M+H]^+^ ion were analyzed in the extracted ion chromatograms (EIC).

### Expression and purification of Spt9

The *spt9* sequence was amplified by PCR with gDNA of *S. oceanensis* CNT-854 as a template and primers: spt9-F, 5’-ctgtacttccaatccggatcc**atgaccgacacccggctact**-3’ (annealing sequence bold faced) and spt9-R, 5’-cagtggtggtggtggtggtgctcgag**tcaacacctggcgttgaccg**-3’ using PrimeSTAR HS (Takara). The pET28a-MBP was digested at BamH1 and XhoI sites. The amplified *spt9* sequence and linear pET28a-MBP were assemble using Gibson assembly (New England Biolabs), and *E. coli* DH 10-beta (New England Biolabs) was transformed with Gibson assembly product. Complete nucleotide sequence of *spt9* in the vector was confirmed by Sanger sequencing. The pET28a-MBP-*spt9* was transformed to *E. coli* Rosetta2 (DE3). *E. coli* Rosetta2 cells harboring pET28a-MBP-*spt9* were grown in Terrific Broth (TB) media containing 50 μg/mL kanamycin and 34 μg/mL chloramphenicol at 37°C until an OD600 of approximately 0.5 and then the temperature was lowered to 16°C. After induction of the T7-promotor with 0.5 mM of Isopropyl β-D-1-thiogalactopyranoside (IPTG), the cells were grown overnight at 16°C. The cells were centrifuged and wash with buffer (20 mM Tris-HCl, 500 mM NaCl, 10% (v/v) glycerol, pH 8.0), and then sonicated in the buffer containing 0.1 mM phenylmethylsulfonyl fluoride (PMSF) on ice. The sonicated cells were centrifuged at 15000 *g* at 4°C, and the supernatant was applied to a His-trap Ni-NTA column (5 mL, GE Healthcare) according to the manufacture instruction using an Äktapurifier platform (GE Healthcare). The eluent containing 300 mM imidazole buffer was desalted using PD-10 desalting column (GE Healthcare) with buffer (20 mM Tris-HCl, 250 mM NaCl, 10% (v/v) glycerol, pH 8.0). The eluent fraction was concentrated by ultra-filtration using Centriprep centrifugal filter YM-10 (Merck Millipore). Protein concentration was determined by Bradford assay (Bio-Rad). For the cleavage of the MBP-tag, the Spt9-H-MBP was treated with Tobacco Etch Virus (TEV) protease at 4°C for 16 hr. The mixture was subjected a His-trap Ni-NTA column and the flow though was collected as the tag-free protein containing fraction. Tag-free Spt9 was desalted and concentrated as described above.

### Expression and purification of Spt6

The *spt6* sequence was amplified by PCR with gDNA of *S. oceanensis* CNT-854 as a template and primers: spt6-F, 5’-tgccgcgcggcagccat**atgagcagtgtcgcggcctacgga**-3’ (annealing sequence bold faced) and spt6-R, 5’-ctcgagtgcggccgcaagctt**gctcatgacgctgccacggtg**-3’ using PrimeSTAR HS (Takara). The pET28a was digested at NdeI and HindIII sites. The amplified *spt6* sequence and linear pET28a were assemble using Gibson assembly (New England Biolabs), and *E. coli* DH10-beta (New England Biolabs) was transformed with Gibson assembly product. Complete nucleotide sequence of *spt6* in the vector was confirmed by Sanger sequencing. The pET28a-*spt6* was transformed to *E. coli* Rosetta2 (DE3). *E. coli* Rosetta2 cells harboring pET28a-*spt6* were grown in TB media containing 50 μg/mL kanamycin and 34 μg/mL chloramphenicol at 37°C until an OD600 of approximately 0.4, and then the temperature was lowered to 18°C. After induction of the T7-promotor with 1.0 mM of Isopropyl β-D-1-thiogalactopyranoside (IPTG), the cells were grown overnight at 18°C. The cells were centrifuged and wash with buffer (20 mM Tris-HCl, 500 mM NaCl, 10% (v/v) glycerol, pH 8.0), and then sonicated in the same buffer on ice. The sonicated cells were centrifuged at 15000 *g* at 4°C, and the supernatant was applied to a His-trap Ni-NTA column (5 mL, GE Healthcare) according to the manufacture instruction using an Äktapurifier platform (GE Healthcare). The eluent containing 300 mM imidazole buffer was desalted using PD-10 desalting column (GE Healthcare) with buffer (20 mM Tris-HCl, 250 mM NaCl, 10% (v/v) glycerol, pH 8.0). The eluent fraction was concentrated by ultra-filtration using Centriprep centrifugal filter YM-10 (Merck Millipore). Protein concentration was determined by Bradford assay (Bio-Rad).

### Enzymatic reaction of Spt9

The purified Spt9-H-MBP and tag-free Spt9 were used for the enzymatic reaction. The reaction mixture containing 100 μM of Spt9-H-MBP, 1 mM *n*-butyl beta-ketoacyl N-acetylcysteamine (**4**) and 1 mM of dihydroxyacetone phosphate (DHAP, **5**) was incubated in 50 mM McIlvaine buffer (pH 7.0) at 30°C for 10 min. 1 volume of MeCN was added to the reaction mixture and then centrifuged at 15000 *g* for 2 min. The filtered supernatant was analyzed by LC-MS using a Phenomenex Luna C18(2) column (5.0 μm, 100 × 4.6 mm, 100 Å) with the following conditions – negative mode, mobile phase A: 0.1% formic acid in H_2_O, mobile phase B: 0.1% formic acid in MeCN, 0.7 mL/min, 0–3 min (10% B), 3–14 min (10%–100% B), and 14–18 min (100% B). The ions at *m/z* 277.0483 (compound **7**) and 279.0639 (compound **8**) corresponding to the [M-H]^−^ ion were analyzed in the extracted ion chromatograms (EIC). The collision energy in the MS/MS analysis of **7** was 10 eV.

### Reduction of butanolide using NaBH_3_CN

50 μL of freshly prepared NaBH_3_CN solution (10 mM in EtOH) was added to the Spt9 enzymatic reaction mixture and incubated at room temperature for 30 min. To this solution, four volumes of 25 mM ammonium acetate buffer (pH 7.6) was added and then applied to the weak anion exchange resin, Macro-prep DEAE (Bio-Rad), packed in a glass pipette (column volume: 0.5 mL). The column washed with buffer (25 mM ammonium acetate buffer, 20% MeOH, pH 7.6), and eluted with 1% NH_4_OH, 20% MeOH aqueous solution. The eluent was filtered and subjected to the LC-MS analysis under the same condition as the Spt9-reaction analysis.

### Reduction of butenolide using *S. tropica* lysate

Lysate was prepared from 25 mL of *S. tropica* CNB-440/Δ*spt9* culture. The mutant was cultured for 4–6 days in A1 media without resin. Cells were harvested from the culture by centrifugation (5000 *g*) and washed with buffer (10 mM Tris-HCl, 330 mM NaCl, 10% (v/v) glycerol, 0.5 mM dithiothreitol (DTT), pH 7.5). Next, the cells were suspended in buffer and sonicated on ice. After removal of cell debris by centrifugation (15000 *g*, 4°C, 10 min), the supernatant was concentrated by ultra-filtration using a Centriprep centrifugal filter YM-10 (Merck Millipore). The resultant solution (protein 15 mg/mL, determined by Bradford assay) was used for the butanolide reduction reaction. This lysate was added to the Spt9 reaction mixture with NADPH (final concentration 1 mM) and then incubated at 30°C for 1 hr. The reaction was stopped by addition of 1 volume of MeCN and then centrifuged at 15000 *g* for 2 min. The supernatant was applied to the LC-MS analysis under the same condition as the Spt9-reaction analysis.

### Dephosphorylation of 8

GBL phosphate (**8**) was incubated with 5 units of calf intestinal alkaline phosphatase (CIP, New England Biolabs) in 50 mM HEPES buffer (pH 8.0) at 30°C for 1 hr. 1 volume of MeCN was added to the reaction mixture and then centrifuged at 15000 *g* for 5 min. The supernatant was applied to the LC-MS analysis after filtration using a spin filter. The lysate was prepared from a 25 mL culture of *S. tropica* CNB-440/Δ*spt9*. LC-MS conditions were the same as Sal-GBLs analysis.

### Enzymatic reaction of Spt6

A 50 μL reaction contained 20 μM Spt6, 1 mM compound **8**, 1 mM ATP, 20 mM MgCl_2_, 50 mM HEPES (pH 8.0). After incubation at 30°C for 60 min, 1 volume of MeCN was added to the reaction mixture and centrifuged at 15000 *g* for 5 min. The supernatant was analyzed by LC-MS/MS in two conditions: (1, Figure 4D, E) TSKgel Amide-80 HR (5 μm, 4.6 × 250 mm, Tosoh), negative mode, mobile phase A: 10 mM ammonium formate H_2_O, mobile phase B: 10 mM ammonium formate in MeCN:H_2_O (9:1, v/v), 0–3 min (90% B, 0.7 mL/min–0.5 mL/min), 3–25 min (90%–50% B, 0.5 mL/min), and 25–30 min (50% B, 0.5 mL/min).(2, Figure S17) Phenomenex Luna C_18_(2) column (5.0 μm, 100 × 4.6 mm, 100 Å), negative mode, mobile phase A: 0.1% formic acid in H_2_O, mobile phase B: 0.1% formic acid in MeCN, 0.7 mL/min, 0–3 min (10% B), 3–14 min (10%–100% B), and 14–18 min (100% B). Ions at *m/z* 279.0639 (compound **8**) and 359.0302 (compound **9**) corresponding to the [M-H]^−^ ion were analyzed in the extracted ion chromatograms (EIC). The collision energy in the MS/MS analysis of **9** was 10 eV.

### Feeding experiments

To a 50 mL preliminary cultured of *S. tropica* CNB-440/Δ*spt9*, isolated Sal-GBL1 (**2**) and synthesized racemic Sal-GBL2 (**3**) in MeOH, or synthesized racemic compound **8** in MeOH were independently added and cultured at 28°C for 1 day with shaking. Autoclaved Amberlite XAD-7HP was added to each culture and fermented for an additional 3 days. The cells and resins were harvested by centrifugation, washed with water, and then extracted with acetone for 2 hr. The extract was filtered and concentrated *in vacuo* using rotary evaporation. After suspension in MeOH, the aliquot was applied to LC-MS for salinipostin analysis as described above.

### Extraction and purification of Sal-GBL1 (2)

*S. tropica* CNB-440 was cultured in 2.8 L Erlenmeyer flasks containing 1 L of medium A1+BFe (A1 plus 1 gL^−1^ CaCO_3_, 40 mgL^−1^ Fe_2_(SO_4_)3·4H_2_O, 100 mgL^−1^ KBr) at 28°C with shaking at 230 RPM (New Brunswick Innova 2300). Sterile Amberlite XAD-7HP resin (20 g, Sigma-Aldrich) was added to a flask after 48 hr from inoculation, and the fermentation was continued for an additional 4 days. The resins and cells from 3 L culture were collected using cheese cloth, washed with water, and soaked in 1.5 L of acetone for 2 hr with gentle shaking. The extract was filtered and concentrated *in vacuo* using rotary evaporation, and the concentrated crude extract was applied to a C_18_ column (Polygoprep C_18_, 25–40 μm, 100 Å (Macherey-Nagel), 40 i.d. × 60 mm). After the column washed with MeOH:H_2_O (1:4, v/v), the compound was eluted with MeOH:H_2_O (1:1, v/v) and MeOH:H_2_O (4:1, v/v) stepwise. The resultant semi-pure extract was purified by HPLC using a Phenomenex Luna C_18_(2) column (5.0 μm, 10 i.d. × 250 mm) with the following conditions – mobile phase A: 0.02% formic acid in H_2_O, mobile phase B: 0.02% formic acid in MeCN, 2.0 mL/min, 0–60 min (50%–100% B). Further purification was performed using a Phenomenex Kinetex C_18_ column (5.0 μm, 10 i.d. × 250 mm) with the following conditions – mobile phase A: 0.02% formic acid in H_2_O, mobile phase B: 0.02% formic acid in MeCN, 2.0 mL/min, 0–10 min (30% B), 10–60 min (30%–100% B). Column fractions were analyzed by MS to obtain purified Sal-GBL1 (**2**, 1.2 mg, clear solid).

### Structure elucidation of Sal-GBL1 (2)

Structure elucidation of Sal-GBL1 (**2**) was accomplished through NMR and CD experiments. The purified compound was dissolved in 0.5 mL of CDCl3. ^1^H NMR, a gradient COSY, a gradient HSQC, a gradient HMBC (^3^*J*_CH_ = 8 Hz) were measured in a 5 mm NMR tube. The residual CHCl_3_ signal at 7.26 ppm in the ^1^H NMR spectrum and the ^13^CDCl_3_ signal at 77.16 ppm in the ^13^C NMR spectra were used as internal references. All the proton and carbon signals were assigned based on the 2D NMR experiments (Figures. S20–23, Table S1). Keto-enol tautomerization gave small signal sets in ^1^H NMR spectrum. The CD spectrum of **2** was measured in MeOH (Figure S24); 279 nm (Δ∊ + 0.465), 229 nm (Δε + 0.342). The observed Cotton effect was compared with that of synthetic 3-(*R*)-A-factor in the literature reported by Mori ^33,34^; 283.5 nm (Δε + 0.699), 221.0 nm (Δε + 0.420).

### Isolation of salinipostin B and G

*S. mooreana* CNT-150 was cultured in 2.8 L Erlenmeyer flasks containing 1 L of medium A1+BFe (A1 plus 1 gL^−1^ CaCO_3_, 40 mgL^−1^ Fe_2_(SO_4_)3·4H_2_O, 100 mgL^−1^ KBr) at 28°C with shaking at 230 RPM (New Brunswick Innova 2300). The cells were harvested from 5 L culture by centrifugation at 15000 *g* for 15 min and soaked in 700 mL of MeOH overnight. After sonication, the crude extract was filtered and concentrated *in vacuo* using rotary evaporation. The concentrate was applied to a C_18_ column (Polygoprep C_18_, 25–40 μm, 100 Å (Macherey-Nagel), 40 i.d. × 60 mm) and eluted step-wise with MeOH:H_2_O (2:3, v/v), MeOH:H_2_O (4:1, v/v) and MeOH. The salinipostin containing fractions were combined and purified by HPLC using a Phenomenex Luna C_18_(2) column (5.0 μm, 10 i.d. × 250 mm) with the following conditions: mobile phase A: 0.02% formic acid in H_2_O, mobile phase B: 0.02% formic acid in MeCN, 2.0 mL/min, 0–60 min (80%–100% B). Salinipostins B and G were identified based on the MS/MS fragmentation pattern, UV profile and ^1^H NMR and COSY spectra^16^.

### Synthesis of beta-ketoheptanoyl-SNAC (4)

As the mimic of beta-ketoacyl-ACP, beta-ketoheptanoyl *N*-acetylcysteamine thioester (**4,** heptanethioic acid, 3-oxo-, *S*-[2-(acetylamino)ethyl] ester) was synthesized ^40,41^. Methyl 3-oxoheptanoate (TCI America, 790 mg, 5 mmol) was dissolved in 6 mL of aqueous 3.0 M NaOH and 300 μL of tetrahydrofuran, and then stirred at room temperature overnight. The residual alcohol was extracted with methylene chloride from the reaction mixture. The aqueous layer was diluted with 50 mL of water and acidified to approximately pH 3 by adding 3 M or 1 M HCl. The acidified mixture was next extracted with 50 mL of methylene chloride. The organic phases were twice washed with saturated aqueous NaCl. The methylene chloride was removed using rotary evaporator, and the resultant clear solid product (424 mg, 59%) was applied to the next reaction without purification. Carboxylic acid (296 mg, 2.1 mmol) was dissolved in the anhydrous methylene chloride (25 mL) and stirred on ice for 15 min. To this solution, *N*-acetylcysteamine (300 mg, 2.5 mmol), 4-(*N*,*N*-dimethylamino) pyridine (DMAP) (63 mg, 0.52 mmol) and *N*-(3-dimethylaminopropyl)-*N*-ethylcarbodiimide hydrochloride (EDCl) (477 mg, 2.5 mmol) were added. This solution was stirred at room temperature overnight. Saturated aqueous NH4Cl solution (25 mL) was added. and the organic phase was collected. The aqueous phase was extracted with ethyl acetate (50 mL) three times. The combined organic phase was concentrated under reduced pressure using rotary evaporation. The residue was purified by silica gel column chromatography with EtOAc to yield **4** (65.1 mg, 13%, clear solid). HR-ESI-MS *m/z* 268.0977 [M+Na]^+^ (calcd for C_11_H_19_NO_3_SNa, 268.0978). ^1^H NMR (CDCl_3_, 600 MHz) δ 3.71 (s, 2H), 3.48 (t, *J* = 6.3 Hz, 2H), 3.11 (t, *J* = 6.3 Hz, 2H), 2.54 (t, *J* = 7.4 Hz, 2H), 2.00 (s, 3H), 1.58 (p, *J* = 7.5 Hz, 2H), 1.35 (m, 2H), 0.92 (t, *J* = 7.4 Hz, 3H); ^13^C NMR (CDCl3, 150 MHz) δ 205.6, 192.6, 170.9, 57.3, 43.3, 39.4, 29.3, 25.6, 23.2, 22.2, 13.9. The signals derived from enol form of **4** were also observed (Figure S27).

### Synthesis and purification of dihydroxyacetone phosphate (5)

Dihydroxyacetone phosphate was synthesized according to the literature ^35^. Ethanol (83 mL) treated with molecular sieves 3A, sulfuric acid (0.5 mL, 9.3 mmol) and triethyl orthoformate (17.5 mL, 105 mmol) were added to a dried round-bottom flask, and refluxed for 30 min under Argon gas. This solution cooled to 4°C, then dihydroxy acetone dimer (2.1 g, 11.7 mmol) was added. After 18 hr, dihydroxy acetone dimer (2.1 g, 11.7 mmol) was added again and stirred for additional 24 hr. Sodium bicarbonate (3.2 g) was added to this solution and stirred at 4°C for 30 min, and then warmed to room temperature. Reaction mixture was filtered through a 2.5 cm of Celite and silica-gel (1:1), and washed with anhydrous ethyl acetate (30 mL). Combined filtrate was concentrated in vacuo and resuspended with ethyl acetate (18 mL) and then evaporated. Hexane (100 mL) was added to the concentrate, and the formed white solid was collected by filtration. The white solid was washed with hexane and dried in vacuum to obtain pure 2,5-diethoxy-*p*-dioxane-2,5-dimethanol (5.37 g, 87 %). Resultant compound (2 g, 8.5 mmol) was dissolved in the pyridine (9 mL), placed on ice for 30 min, and then diphenyl chlorophosphate (3.5 mL, 17 mmol) was added dropwise over 50 min. The reaction mixture was mixed with methyl tert-butyl ether (30 mL) and diethyl ether (10 mL), and washed with H_2_O, cold 1 M HCl, H_2_O, saturated sodium bicarbonate and H_2_O. The organic layer was evaporated in vacuo and dried in vacuum to afford 2,5-diethoxy-*p*-dioxane-2,5-dimethanol *O*-2,*O*-5-bis-(diphenyl phosphate) (yellow oil, 3.81 g, 70%). This compound (1.1 g) was dissolved in ethanol (75 mL) and stirred with platinum oxide (50 mg) under hydrogen gas for 24 hr. After filtration, the reaction mixture was evaporated and dried in vacuum. The resultant colorless oil was dissolved in H_2_O (5 mL) and stirred at 65°C for 5 hr. The pH of this mixture was adjusted to pH 3.7 by 3 M NaOH and H_2_O. In order to improve the purity, a portion of the synthesized product was purified using a HILIC column, TSKgel Amide-80 HR (5 μm, 4.6 × 250 mm, Tosoh), with CH3CN:H_2_O:formic acid (30:70:0.002, v/v/v) as the mobile phase at a flow rate of 0.7 mL/min. HR-ESI-MS *m/z* 168.9910 [M-H]^−^ (calcd for C_3_H_7_O_6_P, 168.9907). Keto-form of **5**; ^1^H NMR (D2O, 600 MHz) δ 4.54 (d, *J* = 7.8 Hz, 2H), 4.46 (s, 2H); ^13^C NMR (D_2_O, 150 MHz) δ 209.5, 66.6, 63.7. Diol-form **5**; δ 3.79 (d, *J* = 5.6 Hz, 2H), 3.53 (s, 2H); ^13^C NMR (D_2_O, 150 MHz) δ 94.6, 67.9, 65.3.

### Racemic synthesis of Sal-GBL2 (3) and Sal-GBL2-phosphate (8)

Sal-GBL2 (**3**) was synthesized based on a literature procedure^42^ in which pentanoyl chloride was instead used to synthesize the desired pentanoyl-gamma-butyrolactone, Sal-GBL2 (**3**). HR-ESI-MS *m/z* 201.1124 [M+H]^+^ (calcd for C10H17O4, 201.1121). ^1^H NMR (CDCl3, 600 MHz) δ 4.43 (dd, *J* = 9.0, 8.1 Hz, 1H), 4.14 (dd, *J* = 9.0, 6.8 Hz, 1H), 3.68 (m, 2H), 3.23 (m, 1H), 2.97 (dt, *J* = 18.0, 7.5 Hz, 1H), 2.63 (dt, *J* = 18.0, 7.3 Hz, 1H), 1.59 (p, *J* = 7.4 Hz, 2H), 1.33 (q, *J* = 7.4 Hz, 2H), 0.90 (t, *J* = 7.4 Hz, 3H); ^13^C NMR (CDCl3, 150 MHz) δ 203.0, 172.4, 69.3, 61.8, 55.1, 42.3, 39.4, 25.4, 22.2, 13.9. The signals estimated to be derived from enol form of **3** were also observed in ^1^H NMR spectrum (Figure S25). Phosphorylation of Sal-GBL2 (**3**) was accomplished using diphenyl phosphoryl chloride in pyridine and successive deprotection by Adams catalyst under hydrogen gas, yielding compound Sal-GBL2-phosphate (**8**, 4.8 mg). HR-ESI-MS *m/z* 279.0636 [M-H]^−^ (calcd for C10H16O7P, 279.0639). ^1^H NMR (CD3OD, 600 MHz) δ 4.45 (br t, *J* = 8.4 Hz, 1H), 4.19 (br t, *J* = 7.5 Hz, 1H), 4.01 (m, 2H), 3.29 (1H), 2.93 (dt, *J* = 18.0, 7.4 Hz, 1H), 2.68 (dt, *J* = 18.0, 7.4 Hz, 1H), 1.58 (p, *J* = 7.4 Hz, 2H), 1.35 (q, *J* = 7.5 Hz, 2H), 0.93 (t, *J* = 7.3 Hz, 3H); ^13^C NMR (CD3OD, 150 MHz) δ 203.3, 173.2, 70.1, 65.6, 43.0, 39.4, 26.0, 22.8, 13.9. The signals estimated to be derived from enol form of **8** were also observed in ^1^H NMR spectrum (Figure S31).

### Distribution and gene content of salinipostin-like biosynthetic gene clusters

ClusterScout searches were performed to identify salinipostin-like biosynthetic gene clusters in other genomes^43^ using the following Pfam functions: *spt1* Pfam00391, Pfam01326; *spt2* Pfam00501; *spt4* Pfam00550; *spt5* Pfam07993; *spt6* Pfam00334; *spt7* Pfam01040; *spt8* Pfam00296; *spt9* Pfam03756. Independent searches were run with a minimum requirement of either 3, 4, or 5 Pfam matches, a maximum distance of <10,000bp between each Pfam match, and a minimum distance of 1bp from the scaffold edge. The boundaries of each match were extended by a maximum of 10,000bp to help return full biosynthetic operons. For some searches, the *spt9*/*afsA* Pfam was defined as essential. MultiGeneBlast^44^ was also used to query the contiguous salinipostin *spt1-9* gene cluster from *S. tropica* CNB-440 against the NCBI GenBank Bacteria BCT subdivision database. Biosynthetic clusters retrieved from each ClusterScout and MultiGeneBlast search were inspected for each of the 9 *spt* Pfam hooks and biosynthetic gene clusters with similar organization to *spt* in *S. tropica* CNB-440 were saved for further analyses.

## ASSOCIATED CONTENT

### Supporting Information

NMR table of **2**; HRMS spectra for **2** and **3**; MS/MS spectra of **2**, **7** and salinipostins; LC-MS chromatograms of **3**, **8**, **9** and salinipostins; SDS-PAGE of Spt9 and Spt6; NMR spectra for **2**−**5**, **8**(PDF)

## AUTHOR INFORMATION

### Notes

The authors declare no competing interests.

## ACKNOWLEDGMENTS

We thank to Dr. Brendan Duggan from UC San Diego for assistance with NMR measurements. We are grateful to Professor M. Reza Ghadiri and Dr. Luke J. Leman at the Scripps Research Institute for their help in the collection of CD measurements. We also thank to our colleagues Dr. Zachary D. Miles, Dr. Gregory C. A. Amos, Dr. Henrique

R. Machado, Dr. Min Cheol Kim, Dr. Xiaoyu Tang, Dr. Hanna Luhavaya, Dr. Jie Li, Dr. Shaun McKinnie and Dr. Jonathan Chekan for experimental advice.

