## Supporting information for "Expansion of gamma-butyrolactone signaling molecule biosynthesis to phosphotriester natural products"

**Table S1.** NMR spectroscopic data for natural Sal-GBL1 (**2**) in $CDCl_3$ (600 MHz).

**Figure S1.** HR-ESI-MS spectra for compounds (**2** and **3**) in the extract of *Salinispora tropica* CNB-440 wild-type.

**Figure S2.** LC-MS chromatograms for Sal-GBL1 (**2**) in the extract of *Salinispora tropica* CNB-440 wild-type cultured with resin.

**Figure S3.** LC-MS chromatograms for compound **3** in the extract of *Salinispora*.

**Figure S4.** HR-LC-MS chromatograms of salinipostins (*m/z* 445) in the extract of *Salinispora*.

**Figure S5.** LC-MS chromatogram and MS/MS spectra of salinipostins in the extract of *Salinispora tropica* CNB-440 WT.

**Figure S6.** LC-MS chromatogram and MS/MS spectra of salinipostins in the extract of *Salinispora arenicola* CNS-250 WT.

**Figure S7.** LC-MS chromatogram and MS/MS spectra of salinipostins in the extract of *Salinispora mooreana* CNT-150 WT.

**Figure S8.** LC-MS chromatogram and MS/MS spectra of salinipostins in the extract of *Salinispora* sp. RL08-036-SPS-B WT.

**Figure S9.** HR-LC-MS chromatograms of salinipostins in the extract of *S. tropica* CNB-440 WT.

**Figure S10.** LC-MS chromatogram and MS/MS spectra of salinipostins (*m/z* 459) in the extract of *S. tropica* CNB-440 WT.

**Figure S11.** LC-MS analysis of salinipostins in the extract of *Salinispora* supplemented with compounds **2**, **3** and **8**.

**Figure S12.** LC-MS chromatogram and MS/MS spectrum of putative salinipostin (*m/z* 445) in the *S. tropica* CNB-440 Δ*spt9* supplemented with **8**.

**Figure S13.** LC-MS chromatogram and MS/MS spectrum of salinipostin B in the *S. tropica* CNB-440 Δ*spt9* supplemented with **8**.

**Figure S14.** LC-MS chromatogram and MS/MS spectrum of putative salinipostin (*m/z* 473) in the *S. tropica* CNB-440 Δ*spt9* supplemented with **8**.

**Figure S15.** LC-MS/MS analysis of enzymatic product (**7**).

**Figure S16.** Dephosphorylation of Sal-GBL2 phosphate (**8**) using CIP.

**Figure S17.** RP-LC-MS analysis of phosphorylation of Sal-GBL2 phosphate (**8**).

**Figure S18.** SDS-PAGE gel (10 %) of purified MBP-Spt9 and tag-free Spt9.

**Figure S19.** SDS-PAGE gel (10 %) of purified His-Spt6.

**Figure S20.** The $^{1}$H NMR spectrum ($CDCl_3$, 600 MHz) of Sal-GBL1 (**2**).

**Figure S21.** The COSY spectrum ($CDCl_3$, 600 MHz) of Sal-GBL1 (**2**).

**Figure S22.** The gradient HSQC spectrum ($CDCl_3$, 600 MHz) of Sal-GBL1 (**2**).

**Figure S23.** The gradient HMBC spectrum ($CDCl_3$, 600 MHz) of Sal-GBL1 (**2**).

**Figure S24.** The circular dichroism (CD) spectrum of Sal-GBL1 (**2**) in MeOH.

**Figure S25.** The $^{1}$H NMR spectrum ($CDCl_3$, 600 MHz) of synthetic Sal-GBL2 (**3**).

**Figure S26.** The $^{13}$C NMR spectrum ($CDCl_3$, 150 MHz) of synthetic Sal-GBL2 (**3**).

**Figure S27.** The $^{1}$H NMR spectrum ($CDCl_3$, 600 MHz) of **4**.

**Figure S28.** The $^{13}$C NMR spectrum ($CDCl_3$, 150 MHz) of **4**.

**Figure S29.** The $^{1}$H NMR spectrum ($D_2O$, 600 MHz) of dihydroxyacetone phosphate (**5**).

**Figure S30.** The $^{13}C$ NMR spectrum ($D_2O$, 150 MHz) of dihydroxyacetone phosphate (**5**).

**Figure S31.** The $^{1}H$ NMR spectrum ($CD_3OD$, 600 MHz) of synthetic Sal-GBL2 phosphate (**8**).

**Figure S32.** The COSY spectrum ($CD_3OD$, 600 MHz) of synthetic Sal-GBL2 phosphate (**8**).

**Figure S33.** The gradient HSQC spectrum ($CD_3OD$, 600 MHz) of synthetic Sal-GBL2 phosphate (**8**).

**Figure S34.** The gradient HMBC spectrum ($CD_3OD$, 600 MHz) of synthetic Sal-GBL2 phosphate (**8**).

**Table S1.** NMR spectroscopic data for natural Sal-GBL1 (**2**) in $CDCl_3$ (600 MHz).[a]

| | Sal-GBL1 (**2**) | | |
|---|---|---|---|
| position | $\delta_{C,}$ type | $\delta_H$ (*J* in Hz) | gHMBC |
| 1 | 172.0, C | | |
| 2 | 54.6, CH | 3.69, d (7.4) | 1, 2, 3, 4, 5, 6 (overlapped to H5) |
| 3 | 38.6, CH | 3.26, m | 1, 2, 4, 5, 6 |
| 4 | 68.6, $CH_2$ | 4.15, d (9.0, 6.8) | 1, 3, 5 |
| | | 4.44, t (8.6) | 1, 2, 3, 5 |
| 5 | 61.5, $CH_2$ | 3.69, dd (10.6, 5.6) | 1, 2, 3, 4, 5, 6 (overlapped to H5) |
| | | 3.73, dd (10.6, 5.5) | 2, 3, 4 |
| 6 | 202.7, C | | |
| 7 | 40.2, $CH_2$ | 2.65, ddd (17.9, 8.8, 6.1) | 6, 8, 9 |
| 7' | | 3.00, ddd (17.9, 9.1, 6.3) | 6, 8, 9 |
| 8 | 31.7, $CH_2$ | 1.51, m | 6, 7, 9, 10, 11 |
| 9 | 27.2, CH | 1.57, m | 7, 8, 10, 11 |
| 10 | 22.0, $CH_3$ | 0.91, d (6.3) | 8, 9, 11 |
| 11 | 22.0, $CH_3$ | 0.91, d (6.3) | 8, 9, 10 |

[a]The $^1$H (600 MHz) and $^{13}$C (150 MHz) NMR spectra were recorded using $CDCl_3$ as the solvent. The signal of residual $CDCl_3$ (7.26 ppm) in the $^1$H NMR spectrum and that of $^{13}CDCl_3$ (77.16 ppm) in the $^{13}$C NMR spectrum were used as the internal references.

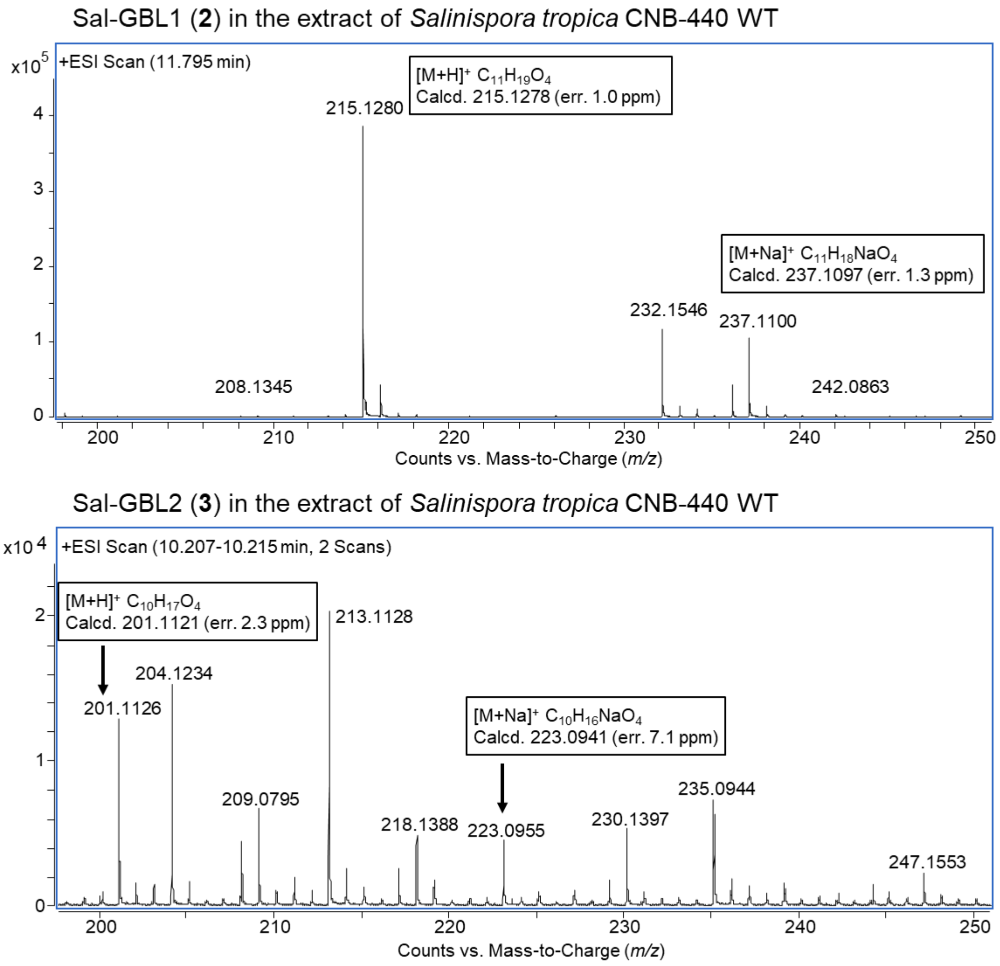


**Figure S1.** HR-ESI-MS spectra for compounds (**2** and **3**) in the extract of *Salinispora tropica* CNB-440 wild-type.

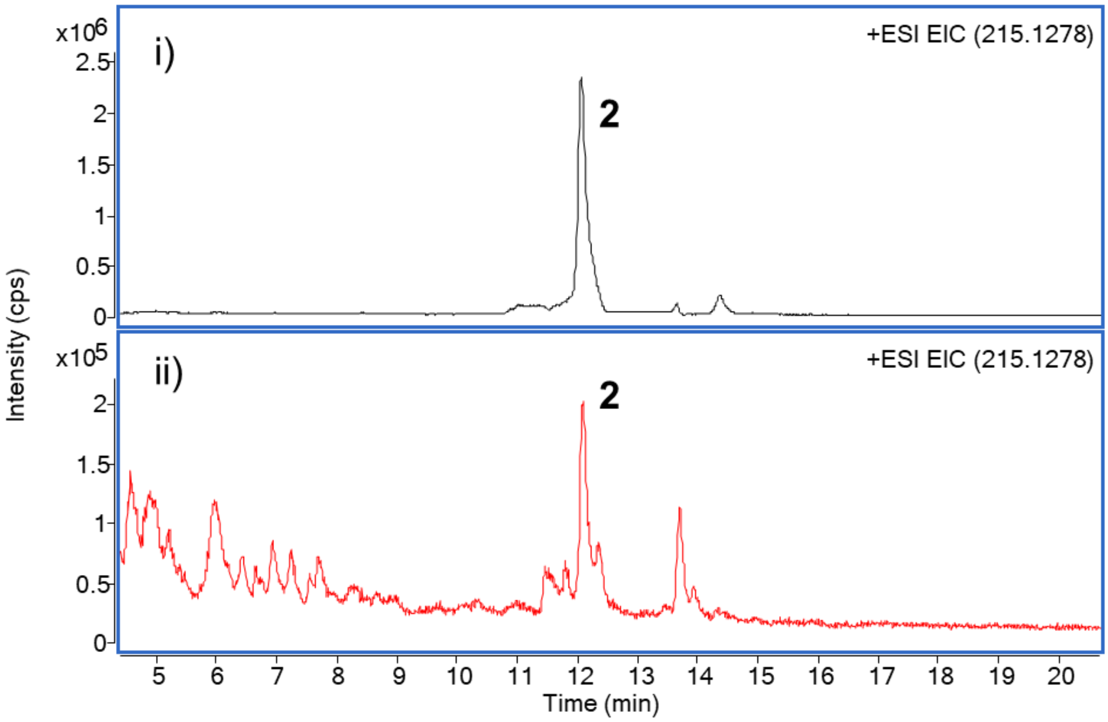


**Figure S2.** LC-MS chromatograms for Sal-GBL1 (**2**) in the extract of *Salinispora tropica* CNB-440 wild-type cultured with resin: i) acetone extract of XAD-7HP and cells, fermented 6 days; ii) ethyl acetate extract from the culture broth fermented 6 days without XAD-7HP. Broth extract (ii) was concentrated 3.5-fold times over the XAD extract (i). Production of **2** was enhanced approximately 54-fold times using resin based on the peak area value. Trace amount of **2** was detected in the cells.

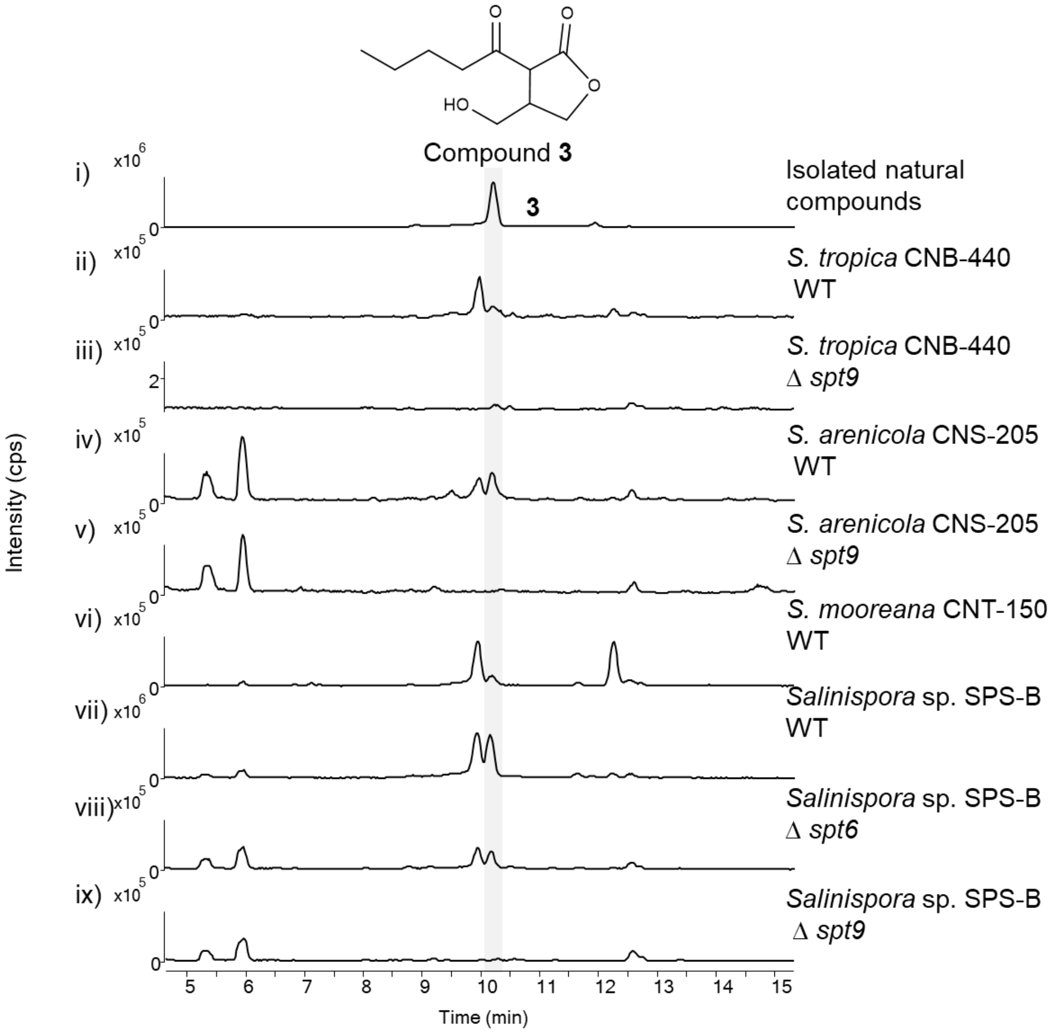


**Figure S3.** LC-MS chromatograms for compound **3** in the extract of *Salinispora*, EIC at *m/z* 201.1121. i) chemically synthesized **3**; ii) *S. tropica* CNB-440 wild type; iii) *S. tropica* CNB-440 Δ*spt9*; iv) *S. arenicola* CNS-205 wild type; v) *S. arenicola* CNS-205 Δ*spt9*; vi) *S. mooreana* CNT-150; vii) *Salinispora* sp. RL08-036-SPS-B WT; viii) *Salinispora* sp. RL08-036-SPS-B Δ*spt6*; and ix) *Salinispora* sp. RL08-036-SPS-B Δ*spt9*.

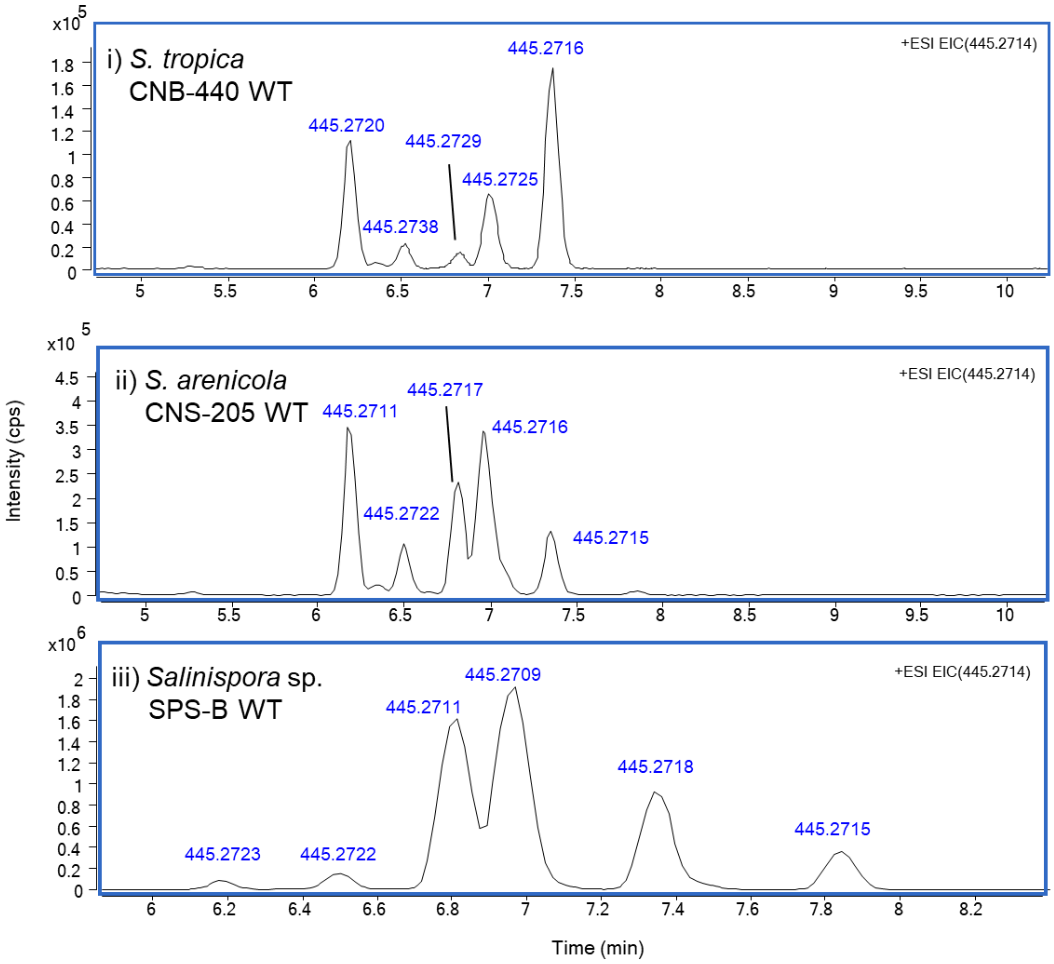


**Figure S4.** HR-LC-MS chromatograms of salinipostins (*m/z* 445) in the extract of *Salinispora*. i) *S. tropica* CNB-440 WT; ii) *S. arenicola* CNS-205 WT; iii) *Salinispora* sp. RL08-036-SPS-B WT. EIC at *m/z* 445.2714 $[M+H]^+$ $C_{23}H_{42}O_6P$. High-resolution mass data are displayed at each compound peak maximum.

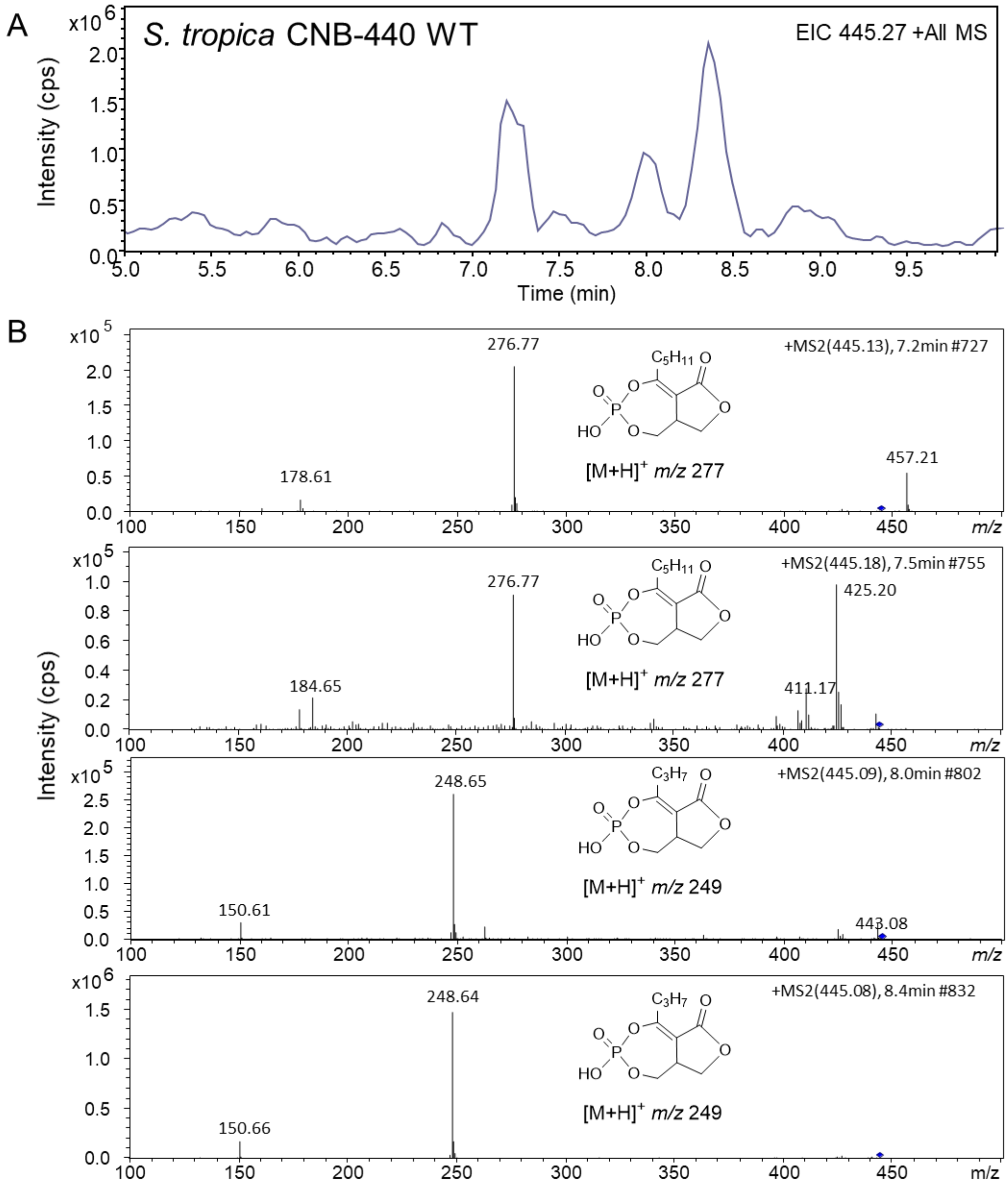


**Figure S5.** LC-MS chromatogram and MS/MS spectra of salinipostins in the extract of *Salinispora tropica* CNB-440 WT. (A) EIC chromatogram at *m/z* 445 (Ion-trap low resolution MS); (B) MS/MS spectra of putative salinipostins and salinipostin G (8.4 min) with the proposed structures of fragment ions.

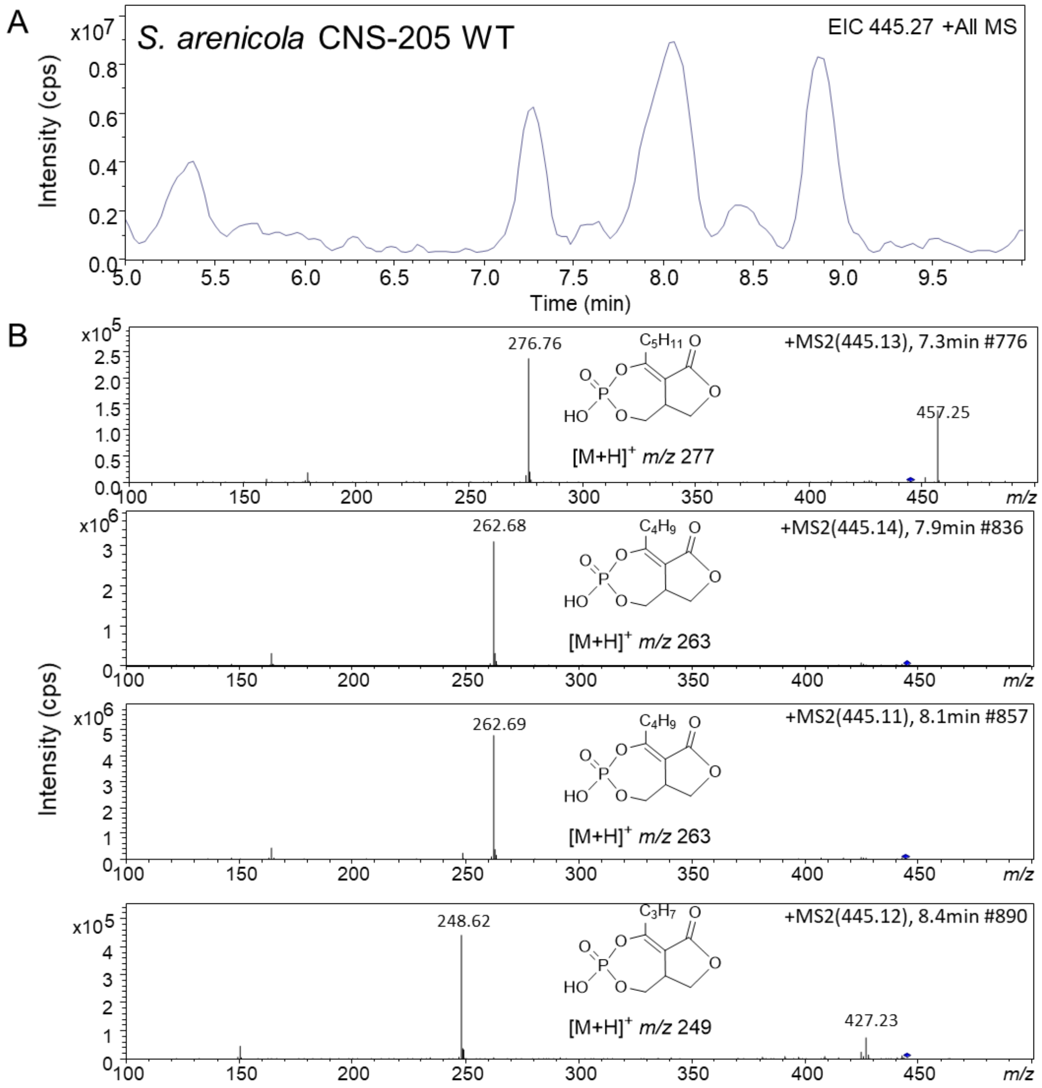


**Figure S6.** LC-MS chromatogram and MS/MS spectra of salinipostins in the extract of *Salinispora arenicola* CNS-250 WT. (A) EIC chromatogram at *m/z* 445 (Ion-trap low resolution MS); (B) MS/MS spectra of putative salinipostins and salinipostin G (8.4 min) with the proposed structures of fragment ions.

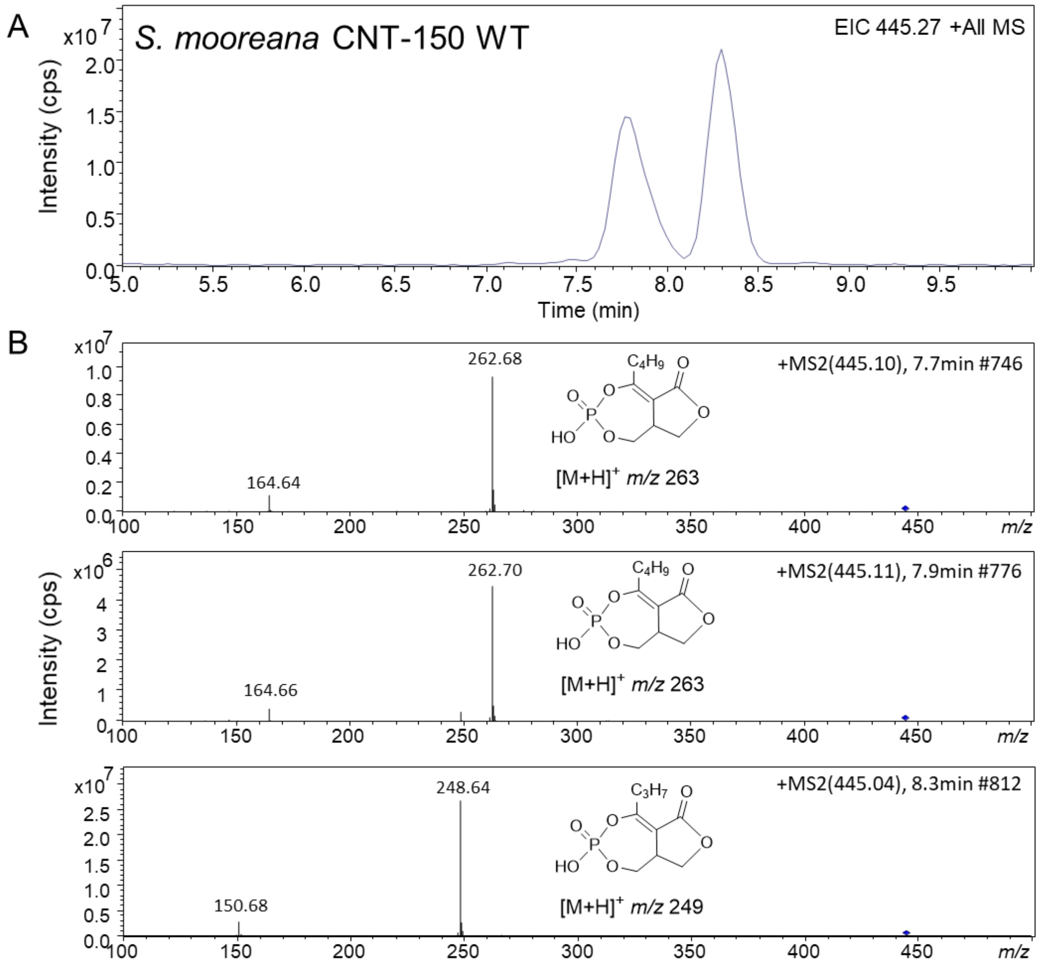


**Figure S7.** LC-MS chromatogram and MS/MS spectra of salinipostins in the extract of *Salinispora mooreana* CNT-150 WT. (A) EIC chromatogram at *m/z* 445 (Ion-trap low resolution MS); (B) MS/MS spectra of putative salinipostins and salinipostin G (8.3 min) with the proposed structures of fragment ions.

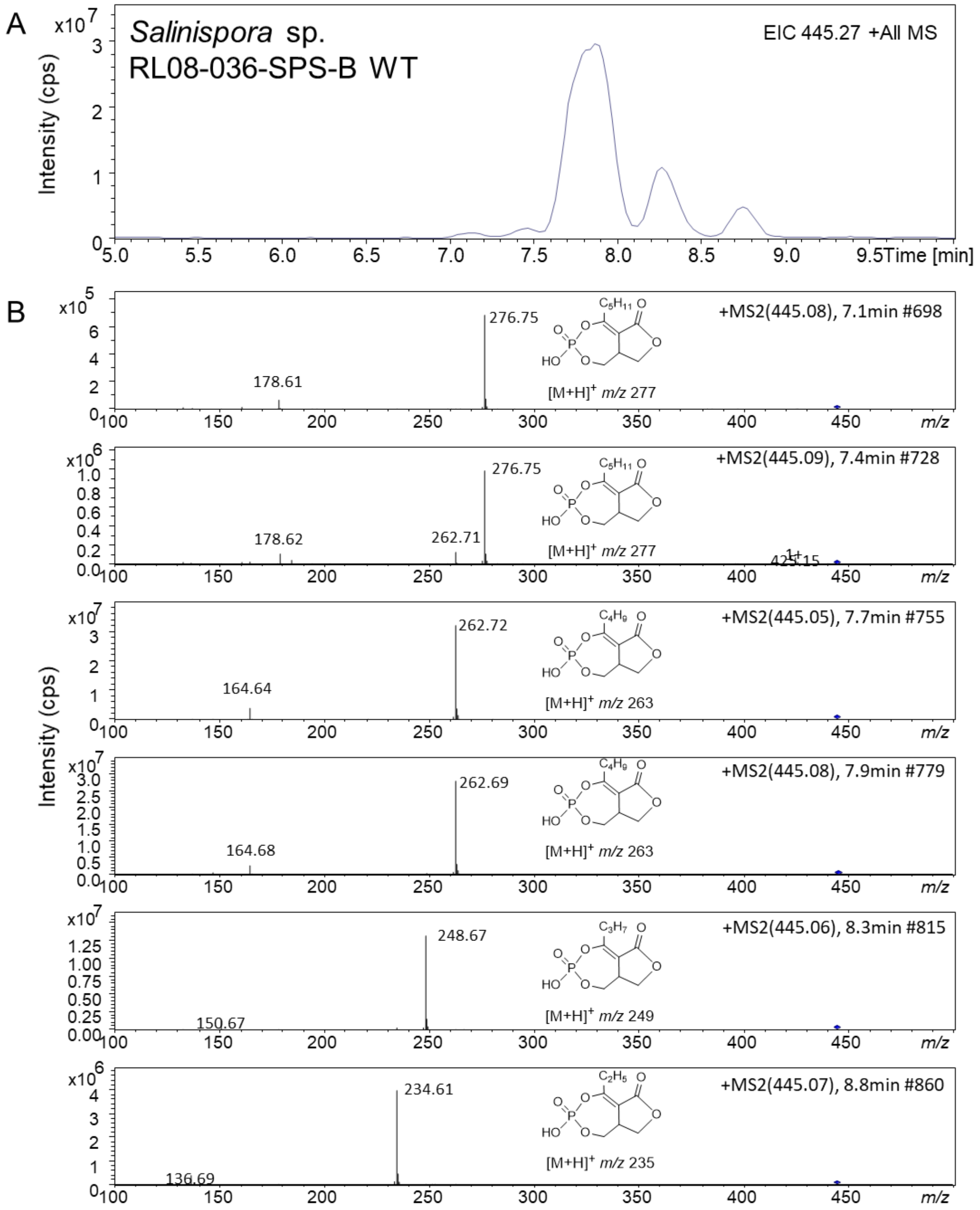


**Figure S8.** LC-MS chromatogram and MS/MS spectra of salinipostins in the extract of *Salinispora* sp. RL08-036-SPS-B WT. (A) EIC chromatogram at *m/z* 445 (Ion-trap low resolution MS); (B) MS/MS spectra of putative salinipostins and salinipostin G (8.3 min) with the proposed structures of fragment ions.

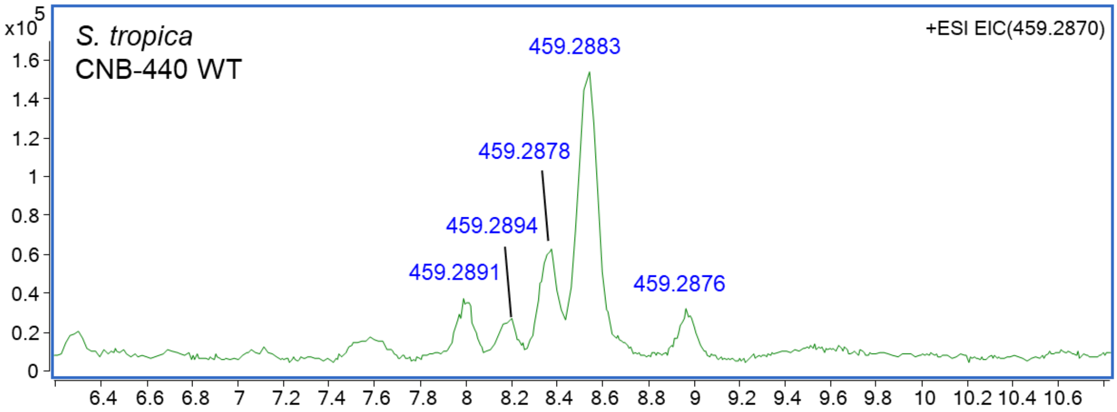


**Figure S9.** HR-LC-MS chromatograms of salinipostins (*m/z* 459) in the extract of *S. tropica* CNB-440 WT. EIC at *m/z* 459.2870 $[M+H]^+$ $C_{24}H_{44}O_6P$. High-resolution mass detected around peak top was displayed.

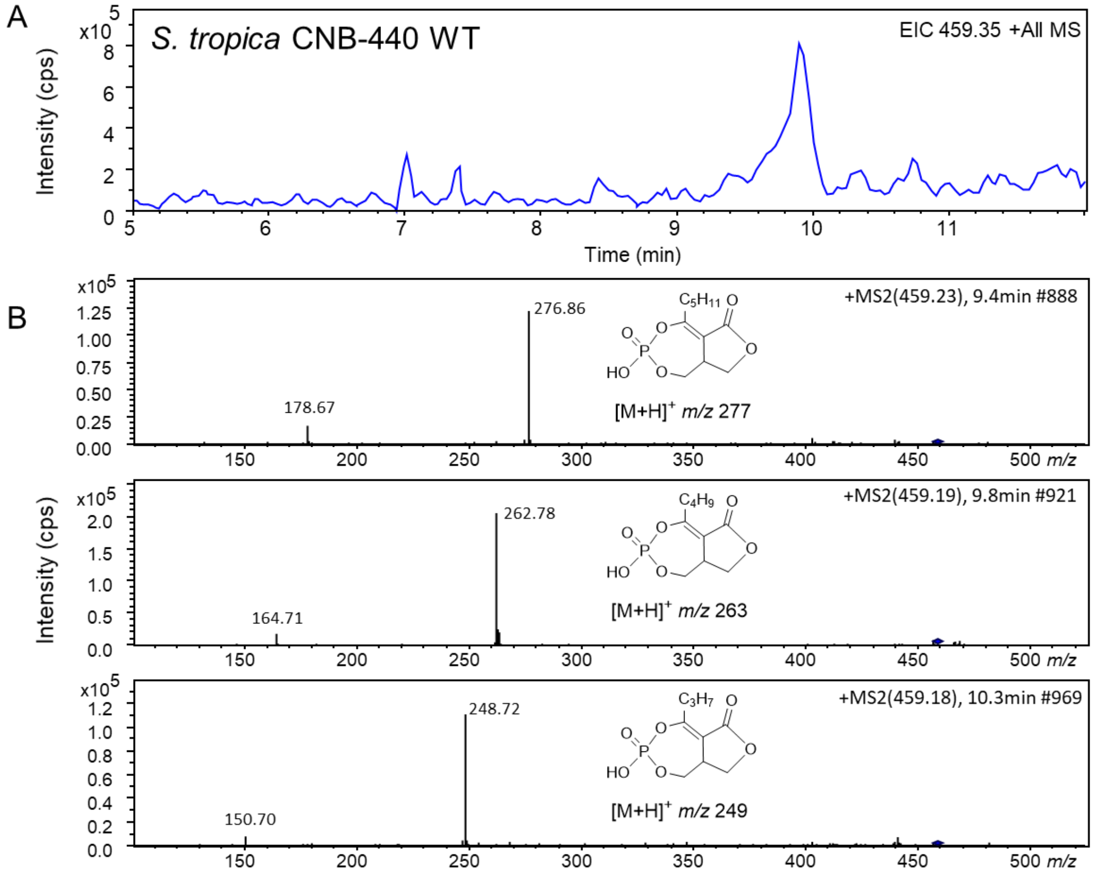


**Figure S10.** LC-MS chromatogram and MS/MS spectra of salinipostins in the extract of *S. tropica* CNB-440 WT. (A) EIC chromatogram at *m/z* 459 (Ion-trap low resolution MS); (B) MS/MS spectra of putative salinipostins and salinipostin B (9.8 min) with the proposed structures of fragment ions.

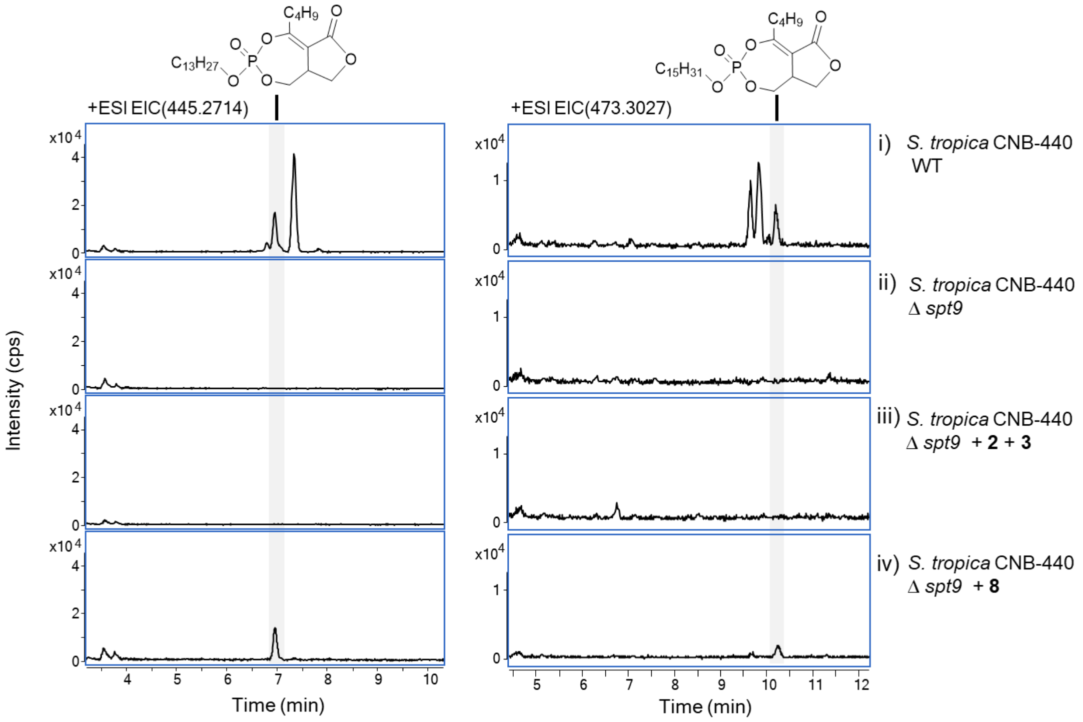


**Figure S11.** LC-MS analysis of salinipostins in the extract of *Salinispora* supplemented with compounds **2**, **3** and **8**, EIC at *m/z* 445.2714 and *m/z* 473.3027. i) *S. tropica* CNB-440 wild type; ii) *S. tropica* CNB-440 Δ*spt9*; iii) *S. tropica* CNB-440 Δ*spt9* supplemented with **2** and **3**; and iv) *S. tropica* CNB-440 Δ*spt9* supplemented with **8**.

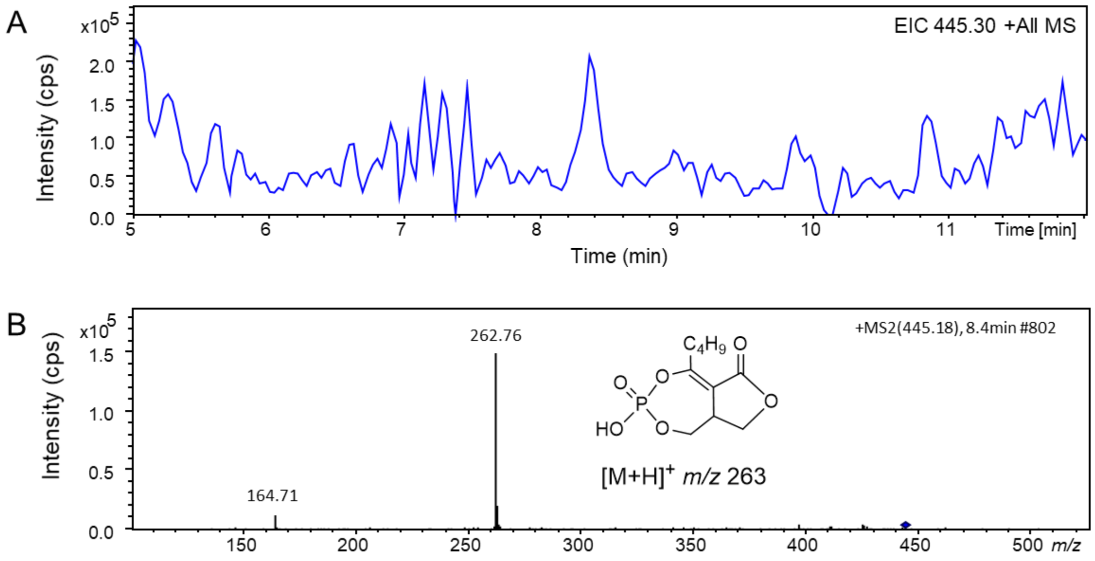


**Figure S12.** LC-MS chromatogram and MS/MS spectrum of putative salinipostin (*m/z* 445) in the *S. tropica* CNB-440 Δ*spt9* supplemented with **8**. (A) EIC chromatogram at *m/z* 445 (Ion-trap low resolution MS); (B) MS/MS spectrum of putative salinipostin with the proposed structures of fragment ions.

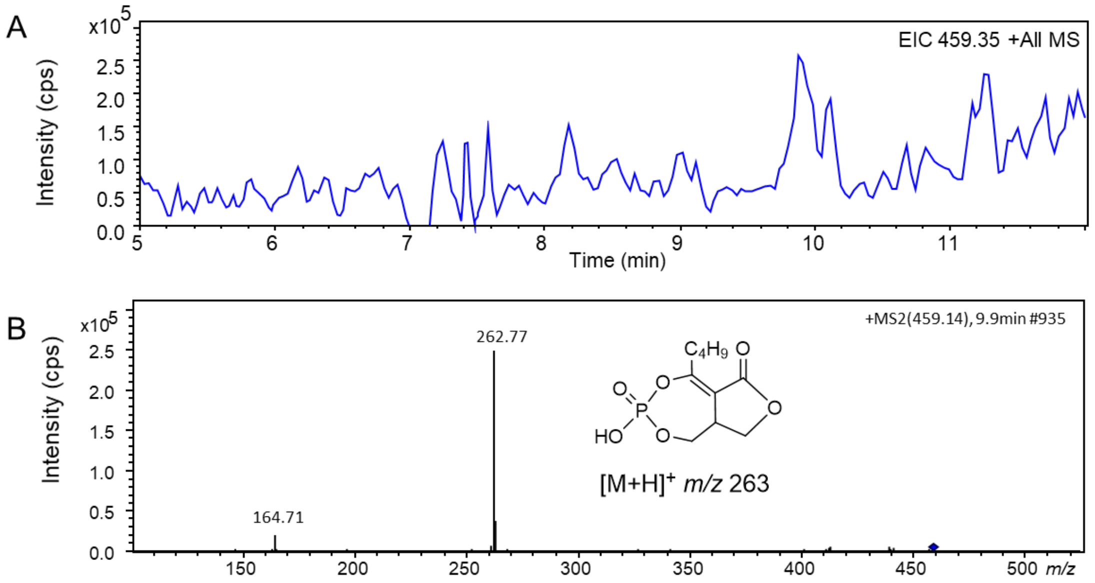


**Figure S13.** LC-MS chromatogram and MS/MS spectrum of salinipostin B in the *S. tropica* CNB-440 Δ*spt9* supplemented with **8**. (A) EIC chromatogram at *m/z* 459 (Ion-trap low resolution MS); (B) MS/MS spectrum of salinipostin B with the proposed structures of fragment ions.

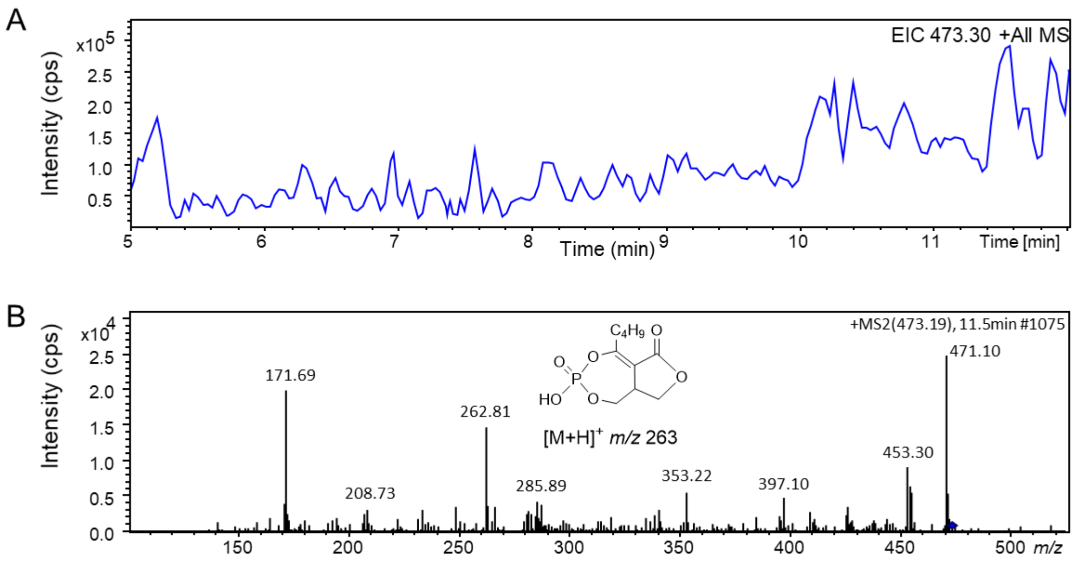


**Figure S14.** LC-MS chromatogram and MS/MS spectrum of putative salinipostin (*m/z* 473) in the *S. tropica* CNB-440 Δ*spt9* supplemented with **8**. (A) EIC chromatogram at *m/z* 459 (Ion-trap low resolution MS); (B) MS/MS spectrum of putative salinipostin with the proposed structures of fragment ions.

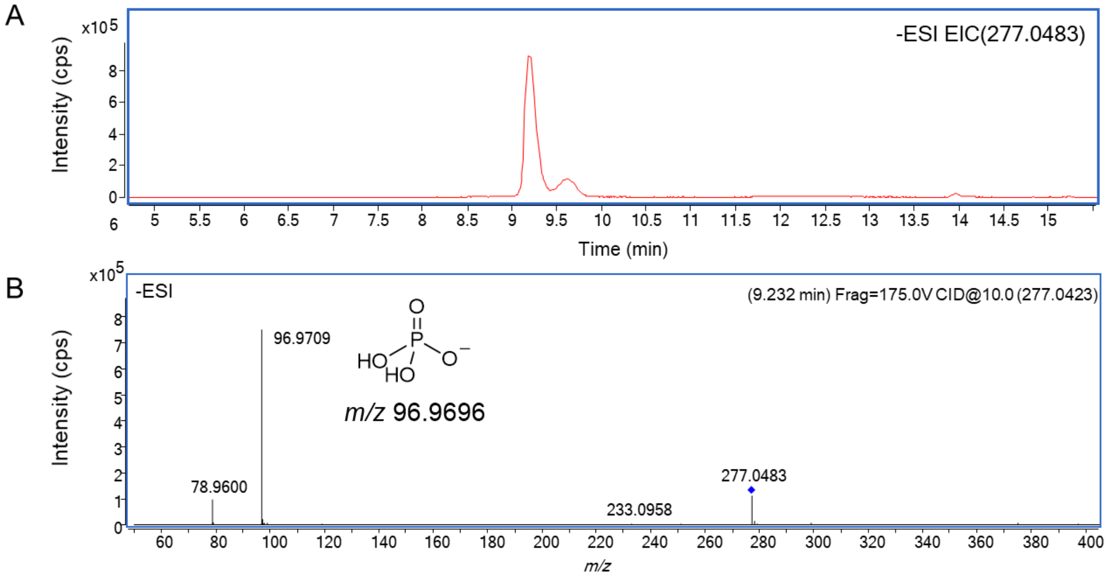


**Figure S15**. LC-MS/MS analysis of enzymatic product (**7**). (A) EIC at *m/z* 277.0483 for **7**; (B) MS/MS spectrum of *m/z* 277. The proposed structure of fragment ion *m/z* 97 was embedded with calculated exact mass.

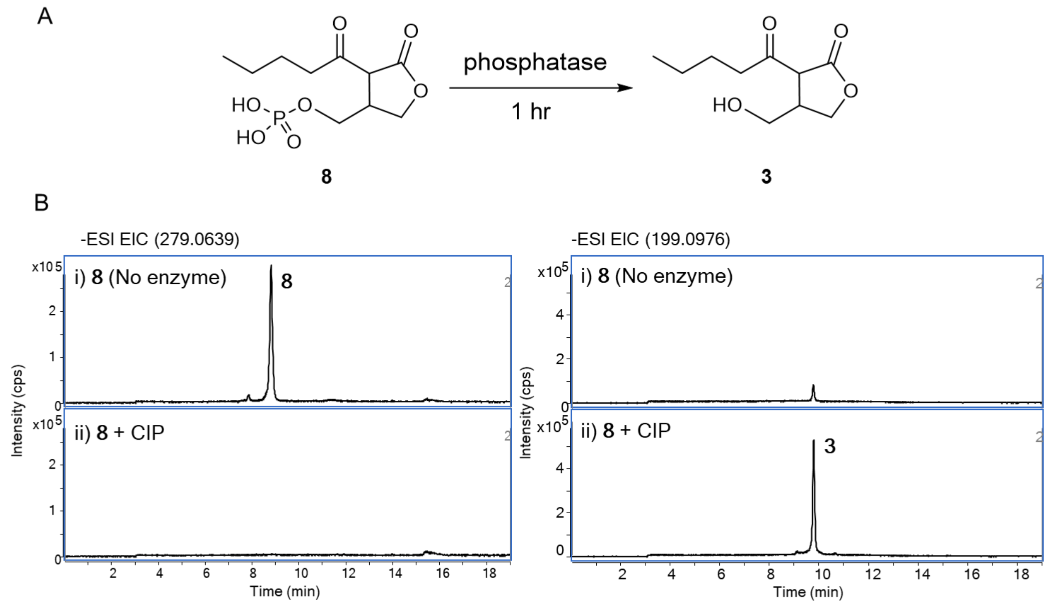


**Figure S16.** Dephosphorylation of Sal-GBL2 phosphate (**8**) using CIP. (A) *In vitro* reaction scheme; (B) Extracted ion chromatograms (EICs) for *in vitro* reaction mixtures: (i) compound **8** without phosphatase; and (ii) **8** with phosphatase (CIP).

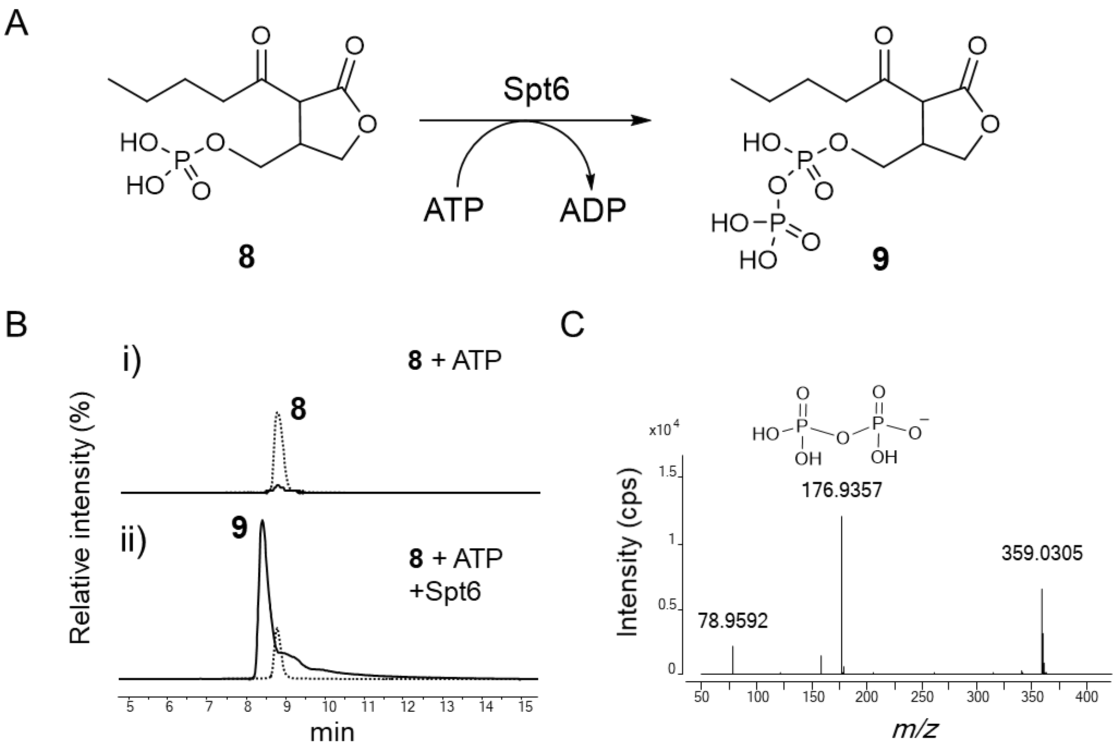


**Figure S17.** RP-LC-MS analysis of phosphorylation of Sal-GBL2 phosphate (**8**). (A) *In vitro* reaction scheme; (B) EICs at *m/z* 279 (dashed line) and *m/z* 359 (plain line): i) **8**, ATP and $Mg^{2+}$ without H-Spt6; and ii) **8**, ATP and $Mg^{2+}$ with H-Spt6. (C) MS/MS spectrum of **9** with the structure of fragment ion, *m/z* 177.

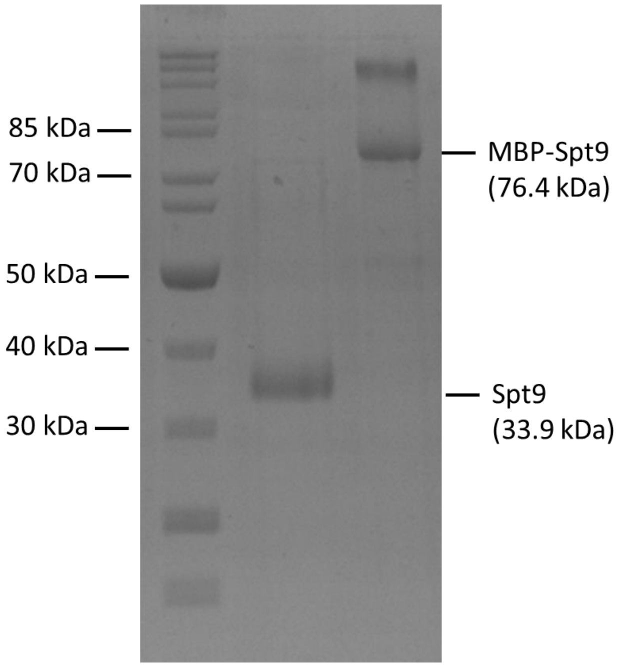


**Figure S18.** SDS-PAGE gel (10 %) of purified MBP-Spt9 and tag-free Spt9.

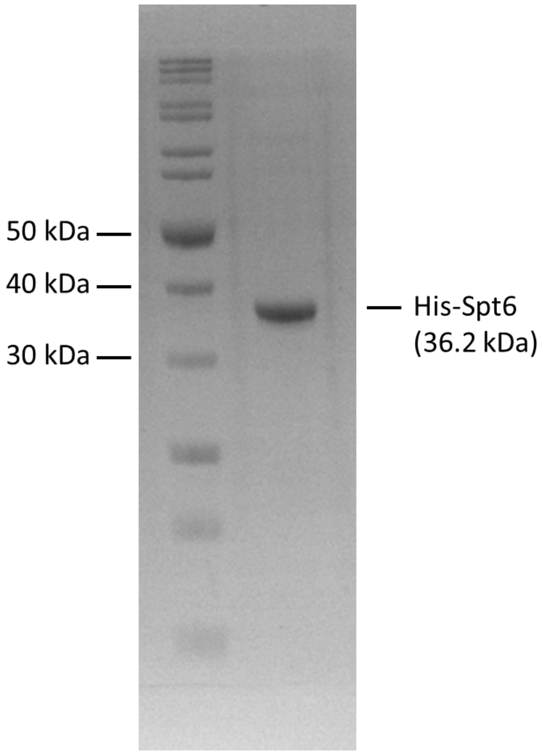


**Figure S19.** SDS-PAGE gel (10 %) of purified His-Spt6.

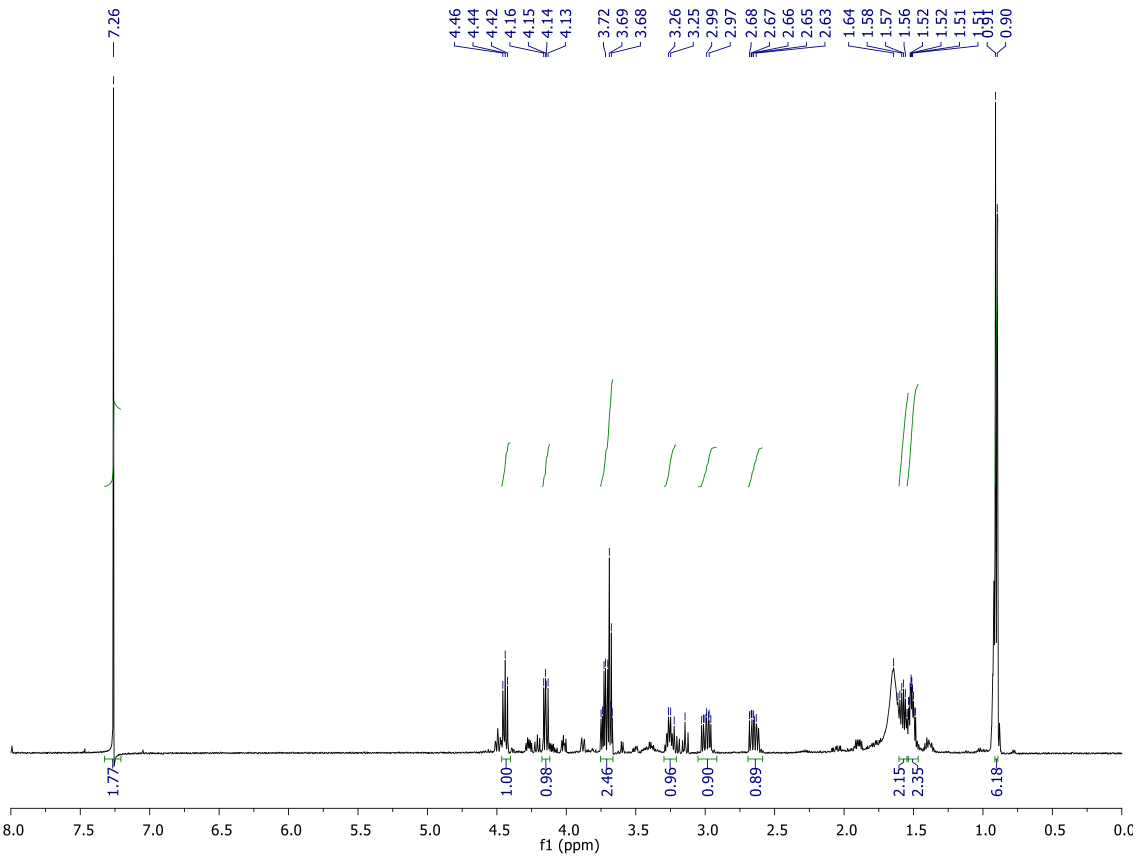


**Figure S20.** The $^{1}H$ NMR spectrum ($CDCl_3$, 600 MHz) of Sal-GBL1 (**2**).

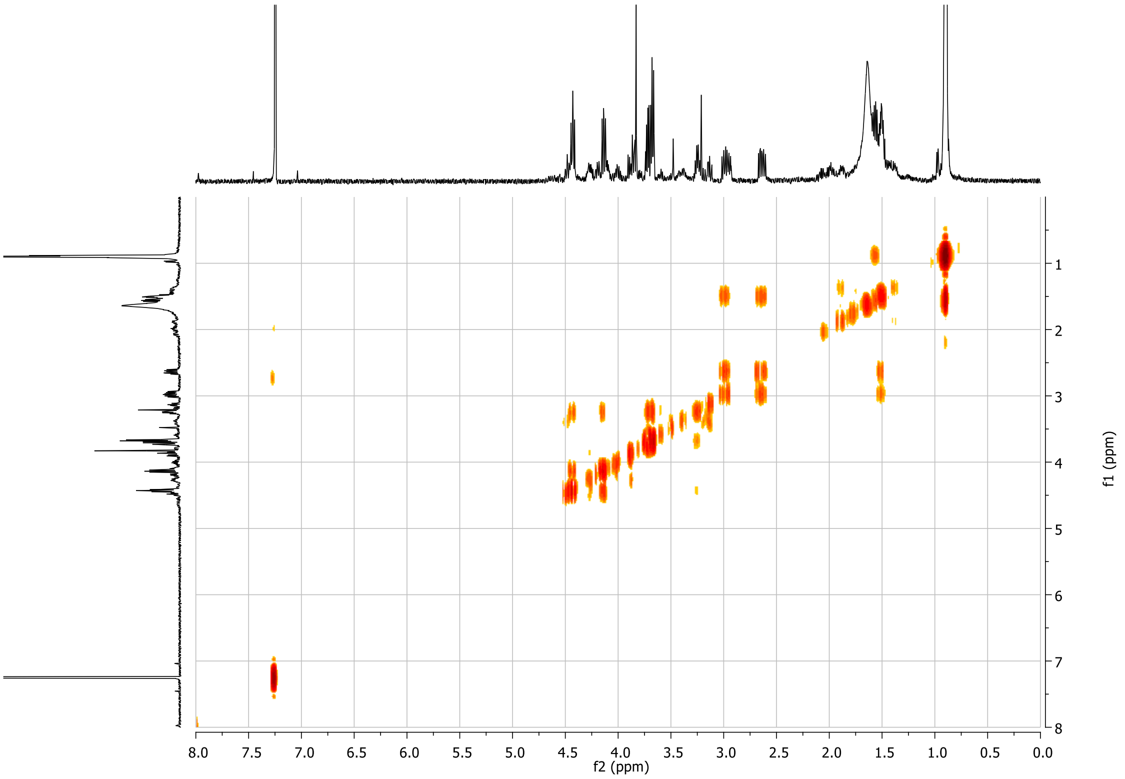


**Figure S21.** The COSY spectrum ($CDCl_3$, 600 MHz) of Sal-GBL1 (**2**).

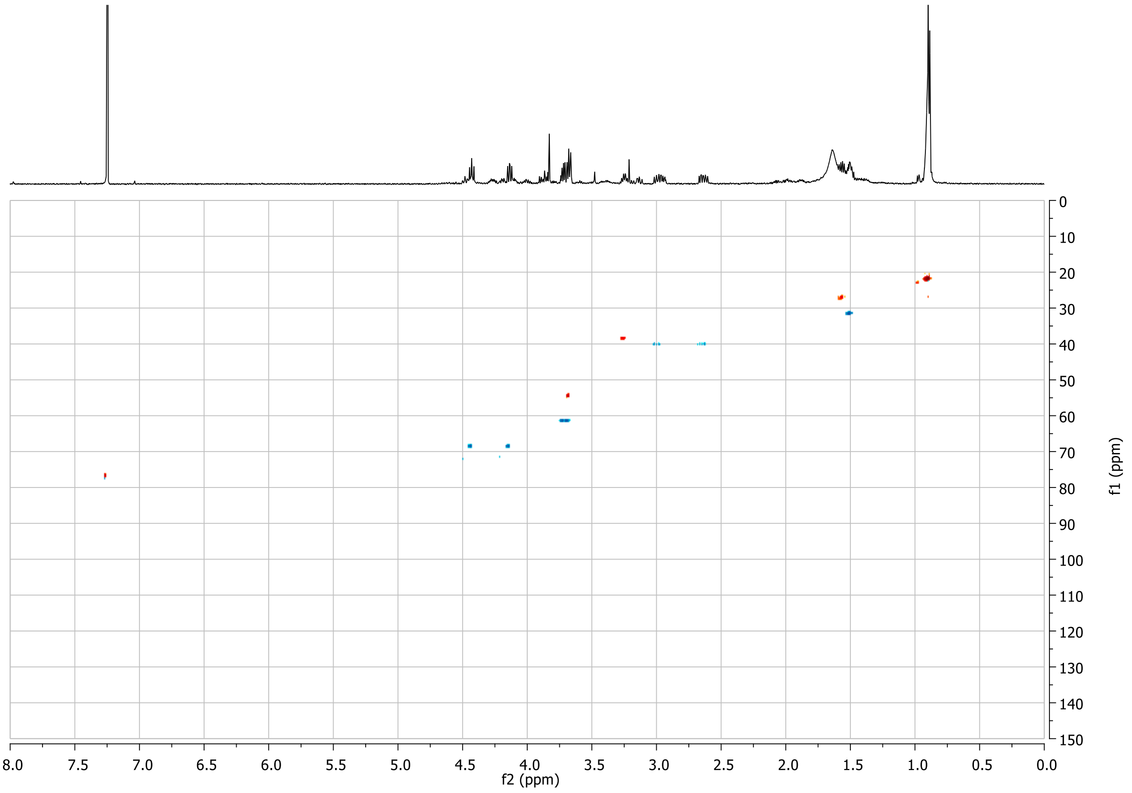


**Figure S22.** The gradient HSQC spectrum ($CDCl_3$, 600 MHz) of Sal-GBL1 (**2**).

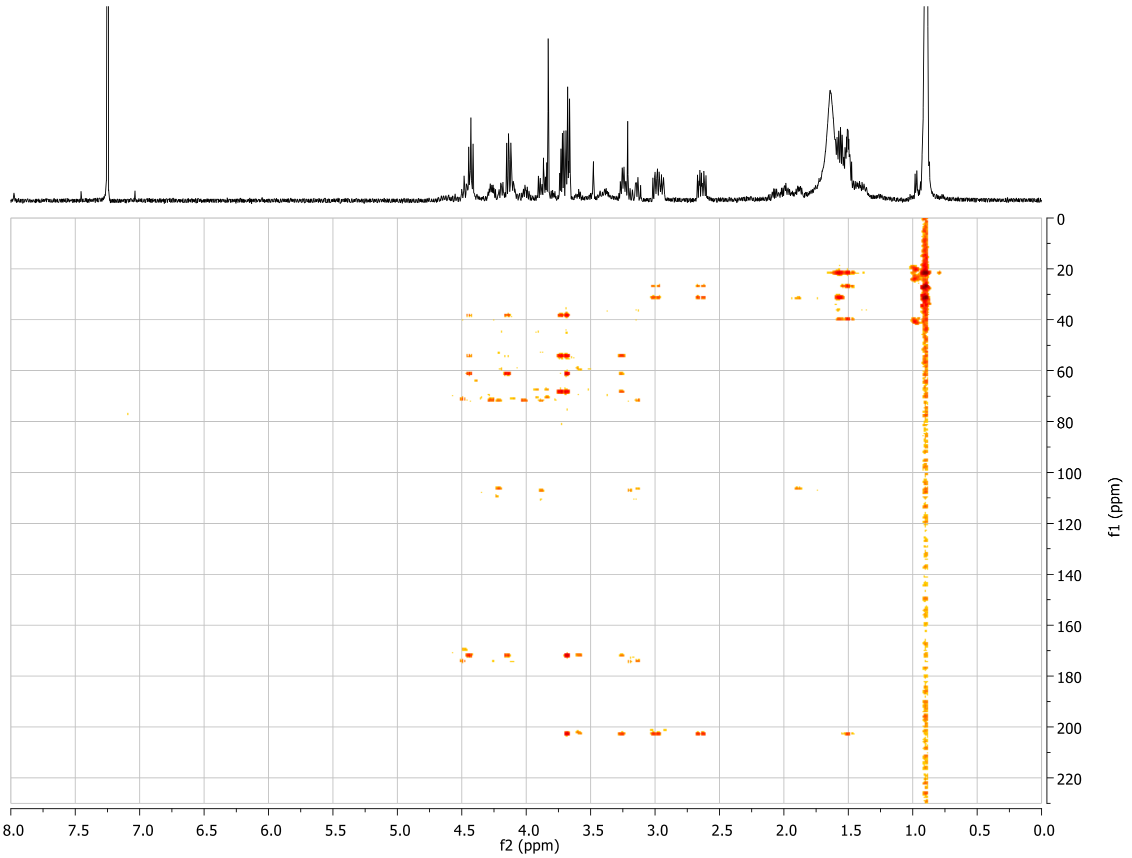


**Figure S23.** The gradient HMBC spectrum ($CDCl_3$, 600 MHz) of Sal-GBL1 (**2**).

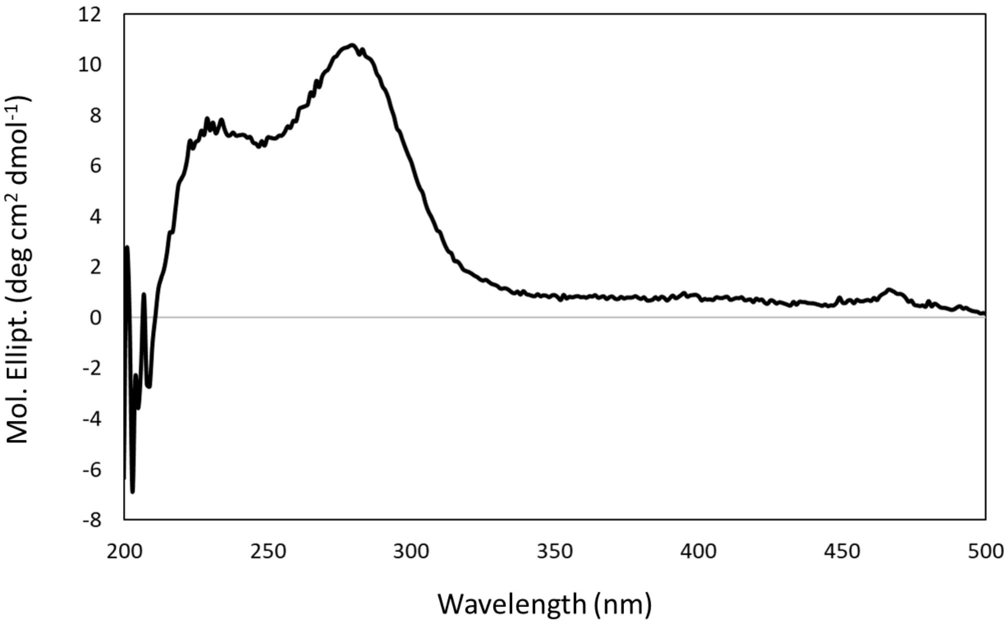


**Figure S24.** The circular dichroism (CD) spectrum of **2** in MeOH.

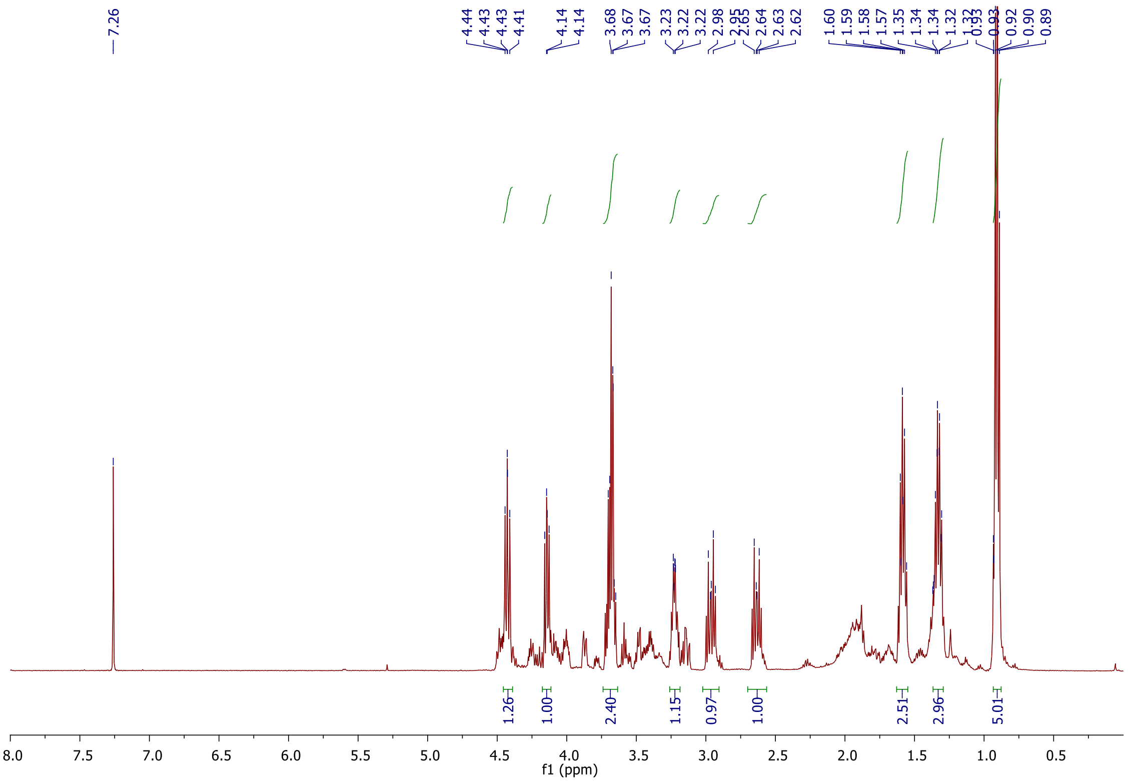


**Figure S25.** The [1]H NMR spectrum ($CDCl_3$, 600 MHz) of synthetic **3**.

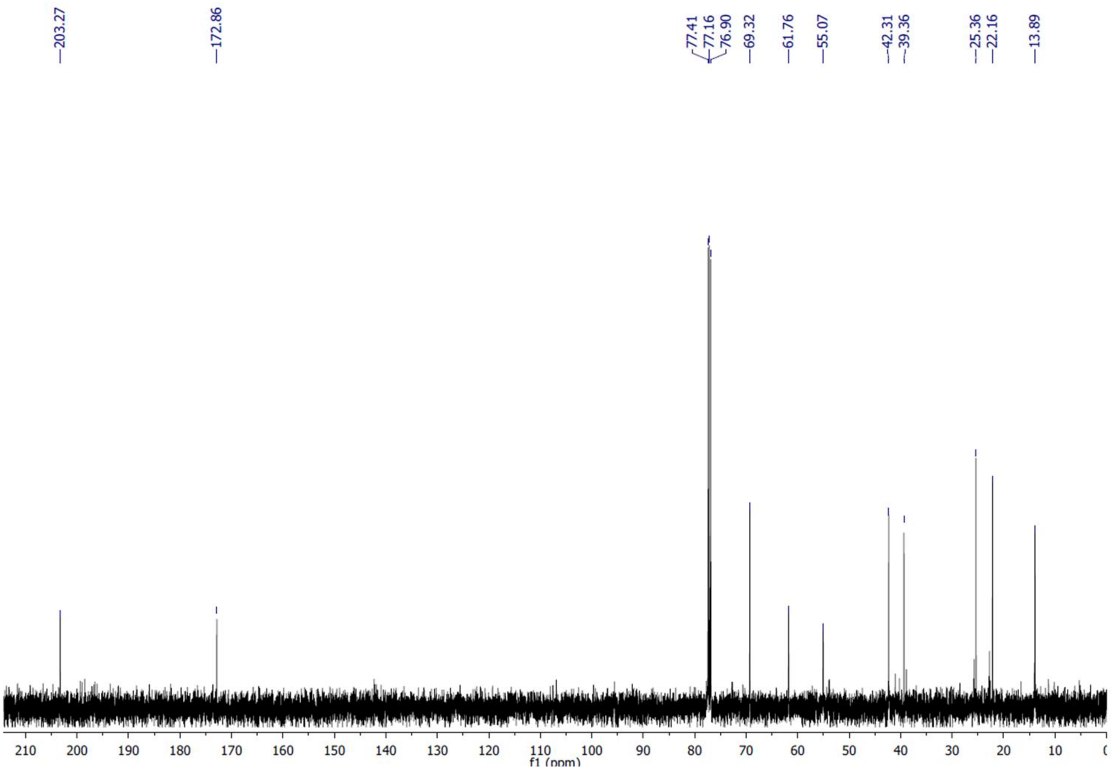


**Figure S26.** The $^{13}C$ NMR spectrum ($CDCl_3$, 150 MHz) of synthetic **3**.

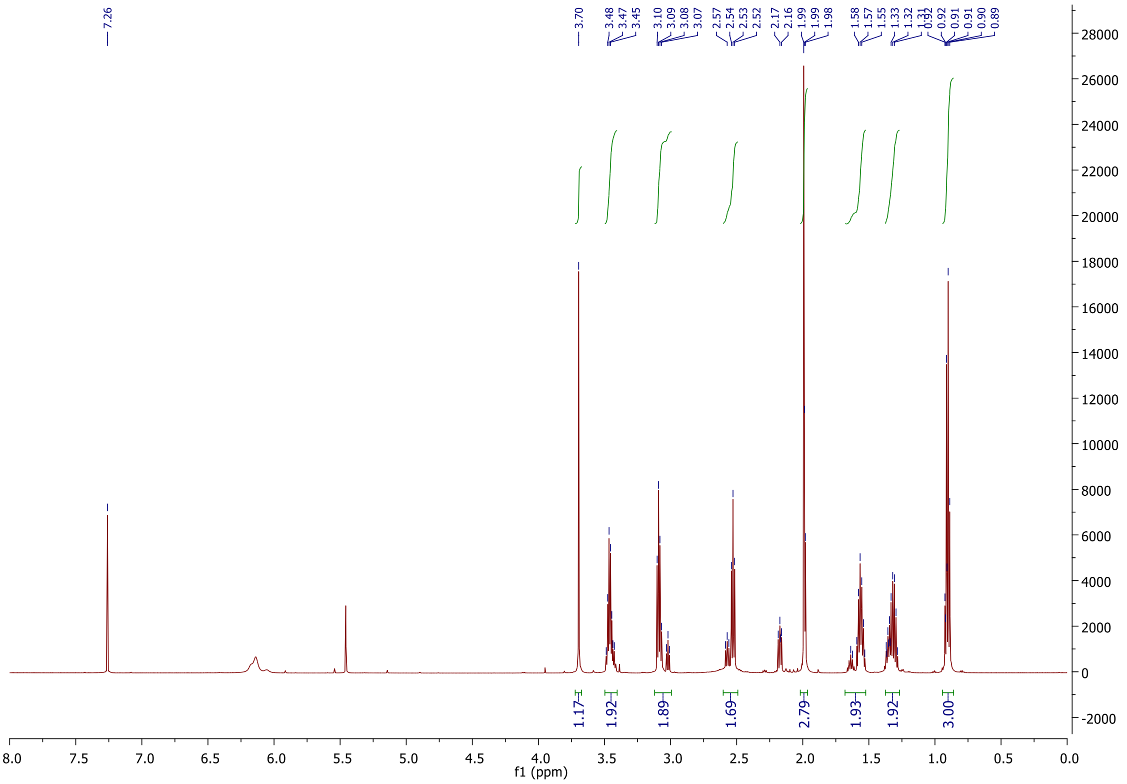

**Figure S27.** The $^{1}H$ NMR spectrum ($CDCl_3$, 600 MHz) of **4**.

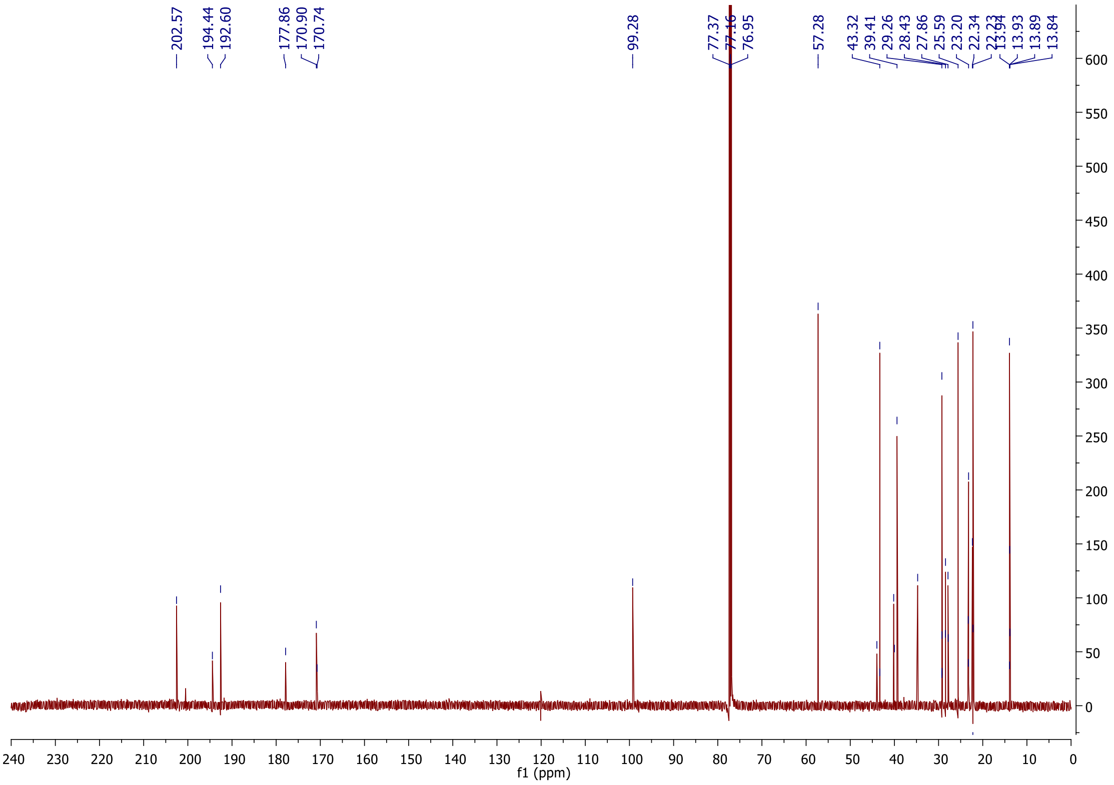


**Figure S28.** The $^{13}C$ NMR spectrum ($CDCl_3$, 150 MHz) of **4**.

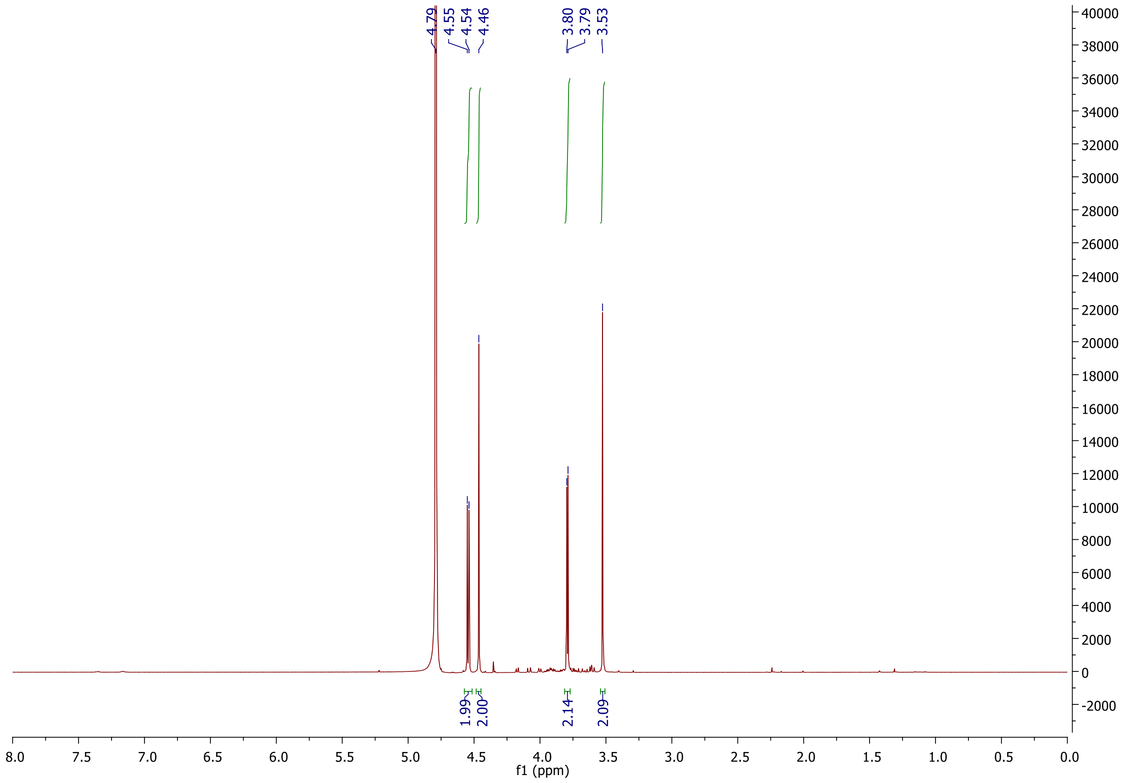


**Figure S29.** The $^{1}H$ NMR spectrum ($D_2O$, 600 MHz) of dihydroxyacetone phosphate (**5**).

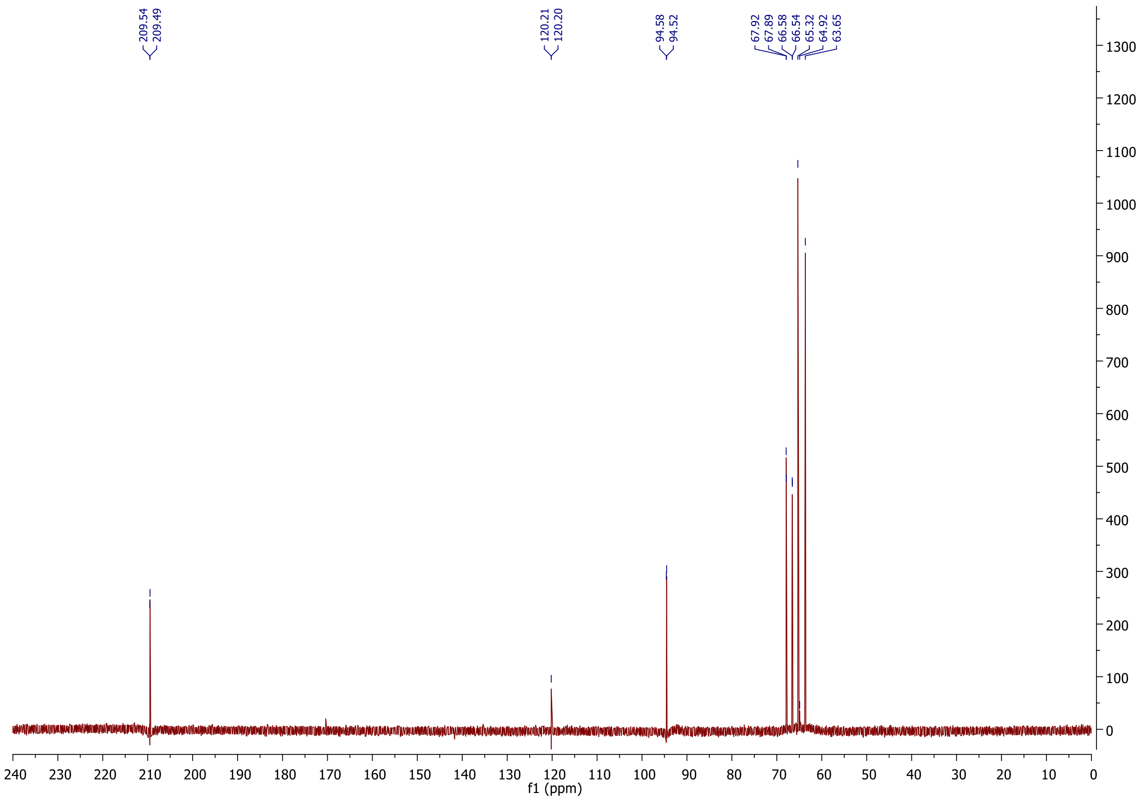


**Figure S30.** The $^{13}C$ NMR spectrum ($D_2O$, 150 MHz) of dihydroxyacetone phosphate (**5**).

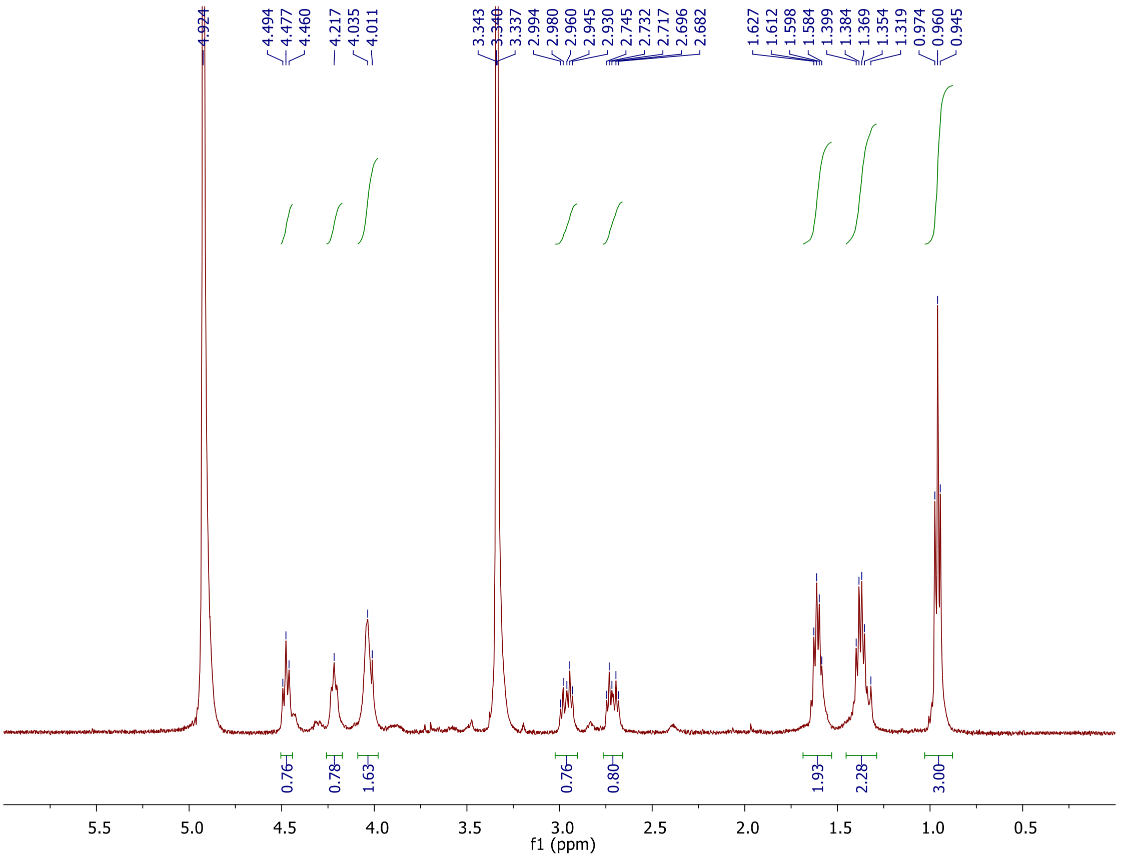


**Figure S31.** The [1]H NMR spectrum ($CD_3OD$, 600 MHz) of synthetic Sal-GBL2 phosphate (**8**).

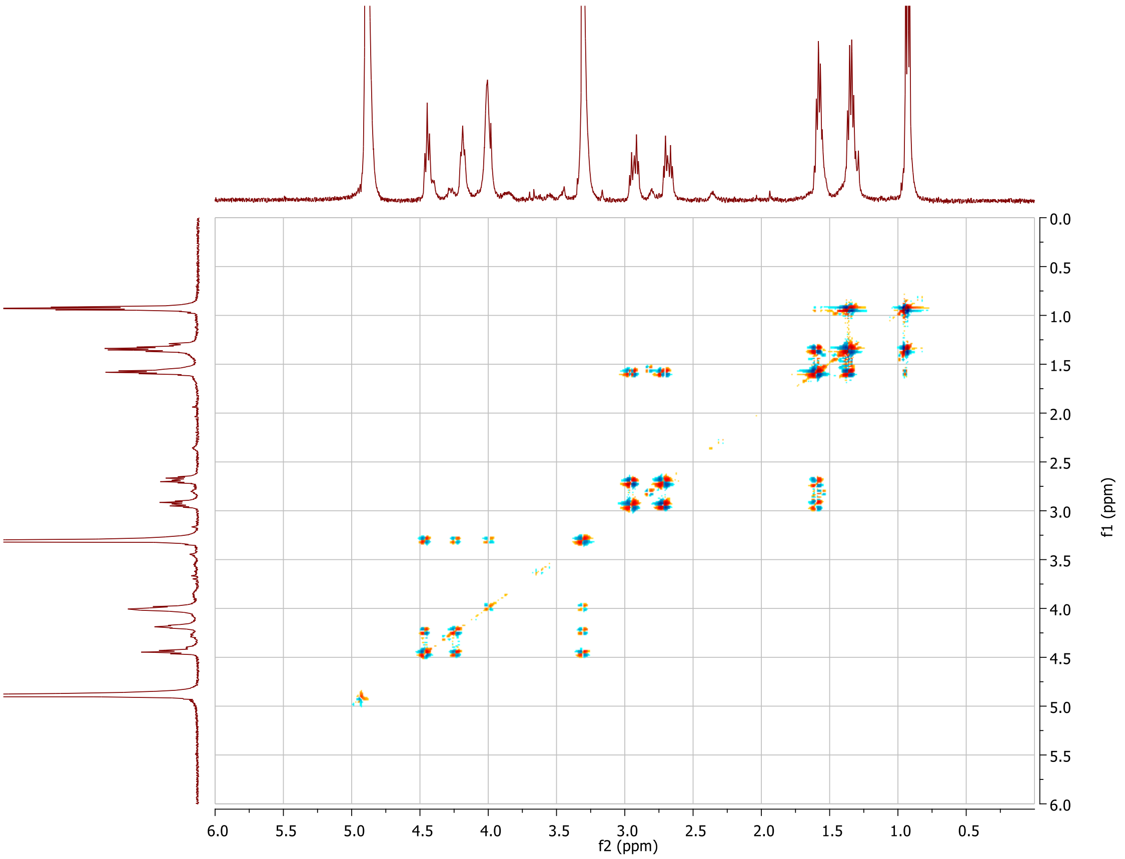


**Figure S32.** The COSY spectrum ($CD_3OD$, 600 MHz) of synthetic Sal-GBL2 phosphate (**8**).

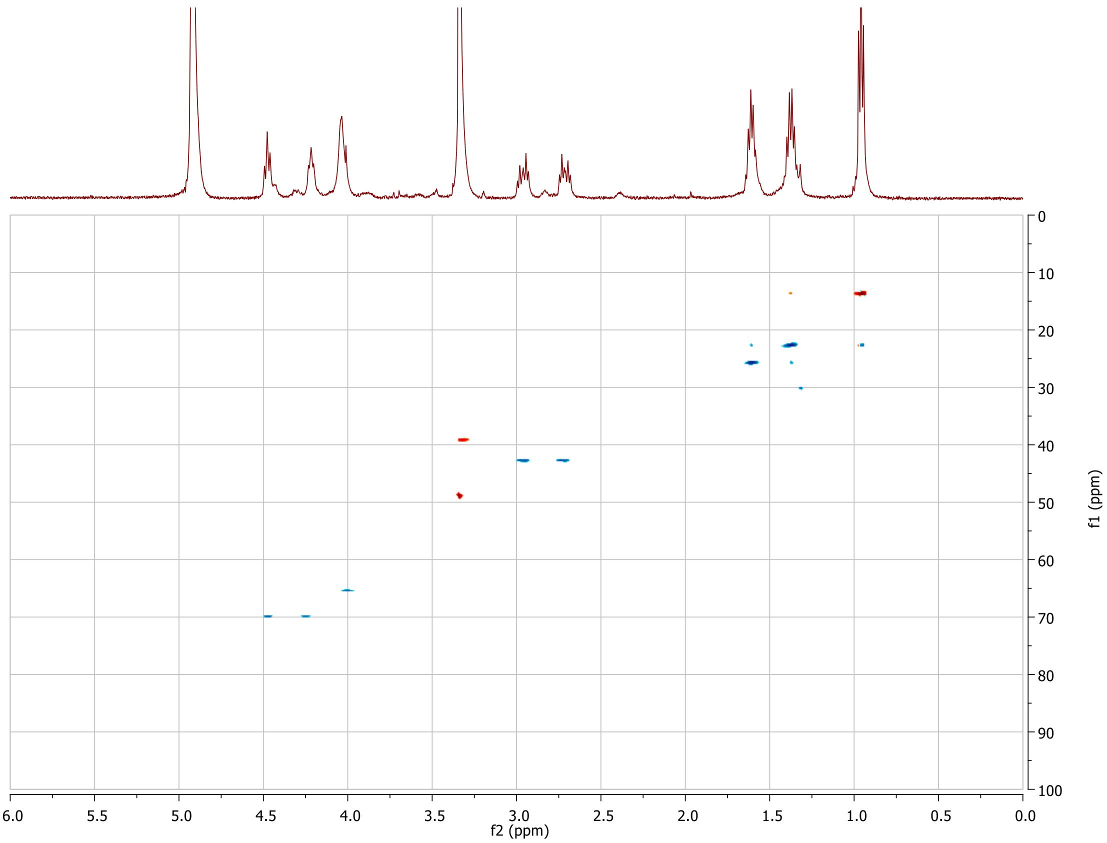


**Figure S33.** The gradient HSQC spectrum ($CD_3OD$, 600 MHz) of synthetic Sal-GBL2 phosphate (**8**).

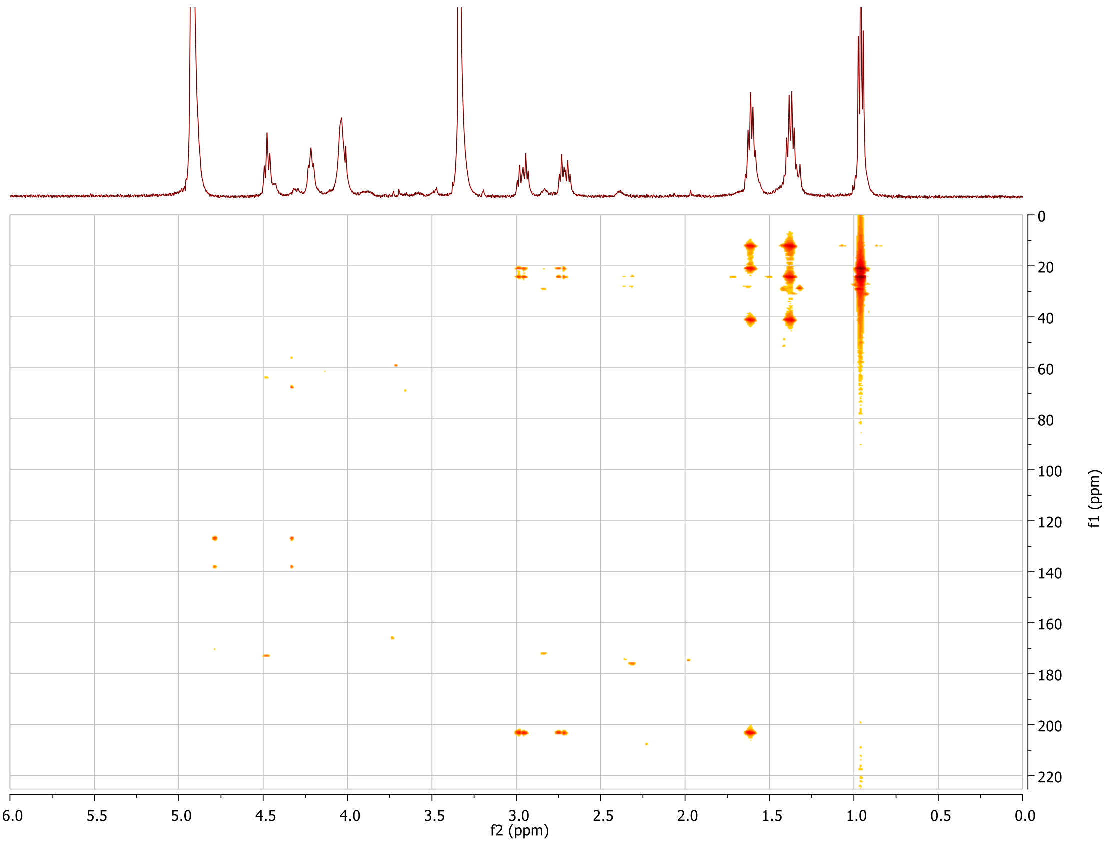


**Figure S34.** The gradient HMBC spectrum ($CD_3OD$, 600 MHz) of synthetic Sal-GBL2 phosphate (**8**).
